# Argonautes and small RNAs associated with nematode programmed DNA elimination

**DOI:** 10.1101/2025.10.23.684123

**Authors:** Maxim V. Zagoskin, Yuanyuan Kang, Richard E. Davis, Jianbin Wang

## Abstract

Programmed DNA elimination (PDE) selectively removes DNA in pre-somatic cells to form the somatic genome during development. Our mechanistic understanding of PDE is mainly based on studies in the single-cell ciliates, where small RNAs and transposases are involved. In contrast, the mechanism(s) of metazoan PDE remains largely unknown. In the parasitic nematode *Ascaris*, PDE removes and remodels all germline chromosome ends and partitions fused germline chromosomes into somatic chromosomes, resulting in dramatic karyotype changes. Here, we identified two *Ascaris* Argonautes, AsWAGO-2 and AsWAGO-3, associated with retained or eliminated chromosome regions, respectively, during PDE mitoses. AsWAGO-3 siRNAs target primarily genes. However, a subset of AsWAGO-3 siRNAs targets repeats within the eliminated regions and located in the middle of chromosomes. These chromosome regions are derived from the fusion of the ends of ancestral chromosomes. In contrast, AsWAGO-2 and its associated siRNAs are transiently enriched on retained chromosomes during elimination mitoses. Overall, our data reveal a link between *Ascaris* WAGOs, their siRNAs, and PDE, identifying small RNA pathways that selectively distinguish retained and eliminated DNA. We suggest that AsWAGO-3 associated small RNAs targeting the interior of select chromosomes may be a response to the fusions of germline chromosomes.

## Introduction

Programmed DNA elimination (PDE) is a striking form of genome remodeling in which specific genomic regions are reproducibly lost ^1,2^. Unlike gradual evolutionary changes or reversible gene regulation via epigenetic mechanisms, PDE imposes a permanent, developmentally programmed alteration to the germline genome to form the somatic genome, challenging the classical notion of genome constancy across cell types ^3,4^. PDE occurs in diverse eukaryotic groups, including ciliates, nematodes, copepods, lampreys, hagfish, anurans, songbirds, and some plants, indicating its biological significance and suggesting its independent evolution in divergent lineages ^5–12^. PDE can be broadly divided into two forms. The first form involves double-strand breaks (DSBs) that lead to chromosome fragmentation, with the broken ends healed by either *de novo* telomere addition or end joining following targeted excision ^5,13^. In the second form, DSB are not required; instead, the entire chromosome(s) are selectively lost during cell division ^10,12^. Studies on single-cell ciliates have provided the most detailed mechanistic insights into the processes of PDE ^14,15^. However, metazoan PDE differs in many respects from ciliate PDE, and its mechanism(s) remain largely unexplored ^16–19^.

In the human and pig parasitic nematode *Ascaris* ^20^, PDE occurs in pre-somatic cells during embryogenesis at the 4-16 cell stages, while the genome in the germline cells remains intact ^5^. Tandem 121-bp satellite repeats and ∼1,000 germline-expressed genes are eliminated, suggesting PDE serves to irreversibly silence repeats and genes in the somatic cells ^21^. During *Ascaris* PDE, DSBs are introduced at ∼100 sites, known as chromosome breakage regions (CBRs) ^17,22,23^. The broken DNA ends are resected and healed with telomere addition ^17,24^. Interestingly, no conserved motif or other sequence features are present in the CBRs, suggesting a sequence-independent mechanism for *Ascaris* PDE ^22^. *Ascaris* has holocentric chromosomes, and its centromeres are reorganized during development and PDE. Regions destined for elimination have reduced centromeres and are not segregated during PDE mitoses ^19^. The eliminated DNA is sequestered into micronuclei, which are subsequently relegated to the cytoplasm for degradation through phagocytosis ^23^.

All ends of the 24 *Ascaris* germline chromosomes are removed, including subtelomeric regions and telomeres ^23^. Interestingly, 12 regions in the middle of 11 germline chromosomes are also eliminated (chromosome X1 has two internal regions removed). These internal breaks appear to occur at sites of ancient fusions of ancestral chromosomes ^25^. These fusions reduce the number of germline chromosomes and could serve to reduce meiotic errors ^25^. Chromosome fusions are exemplified to the extreme in the horse parasite *Parascaris*, which has a single pair of germline chromosomes ^26^. Remarkably, after PDE, the distinct germline genomes of *Ascaris* (24 chromosomes) and *Parascaris* (1 chromosome) are partitioned into the same set of 36 somatic chromosomes, thus resolving the fused chromosomes ^25^. This suggests that fusion and PDE play a crucial role in shaping the genomes of these nematodes ^25^. It also raises questions with respect to the significance of fusions and PDE and whether and to what extent they impact the 3D genome organization, the regulation of gene expression, and the cellular response to these karyotype changes.

Small RNAs play a central role in guiding ciliate PDE, marking retained or eliminated DNA during the development of the somatic macronucleus. In *Paramecium* and *Tetrahymena*, small RNAs (also known as scan RNAs or scnRNAs) associate with PIWI-clade Argonaute proteins to identify sequences targeted for elimination ^14,27^, whereas in *Oxytricha* and *Euplotes*, small RNAs specify genomic regions for retention ^28,29^. These small RNAs and Argonaute complexes mediate chromatin modifications by recruiting PRC2-like complexes that deposit repressive histone marks such as H3K9me3 and H3K27me3 at loci destined for elimination ^11,30,31^. Interestingly, in *Euplotes*, small RNAs were recently suggested to identify the precise boundary of some DNA break sites ^29^. In *Ascaris* early embryos, the histone variant CENP-A is differentially deposited onto the retained but not eliminated DNA to facilitate PDE ^19^. Given the precedence of small RNAs and PDE in ciliates, small RNAs and Argonautes might participate in *Ascaris* PDE through 1) the recognition of eliminated versus retained regions, 2) the identification of DNA break sites, 3) interactions with centromeric or subtelomeric/telomeric chromatin, and/or 4) their silencing of genes and repeats within eliminated regions.

Nematodes have an expanded set of Argonautes and their associated small RNAs ^32^. In *Caenorhabditis elegans*, 19 functional argonautes have been identified and characterized ^33^. They are associated with miRNAs, piRNAs (21U-RNAs), 26G-RNAs, and 22G-RNAs, and they function as regulatory RNAs ^34,35^. *Ascaris* has ten Argonautes associated with miRNAs, 26G-RNAs, and 22G-RNAs ^36,37^. Interestingly, *Ascaris* lacks piRNAs or the PIWI-clade Argonautes, the clade involved in ciliate PDE ^30,38^. However, small RNA pathways are flexible in nematodes, and it is thought that the loss of the piRNAs is compensated by a robust set of endogenous siRNAs expressed during *Ascaris* germline development ^37^.

In a previous study ^37^, we characterized the diversity and developmental dynamics of *Ascaris* Argonautes and their associated small RNAs across the germline and early embryogenesis. While the work included analysis of early embryos undergoing PDE, no clear association between WAGO-bound small RNAs and PDE was detected. However, the previous analysis was performed in the whole embryos and in a global developmental context, where WAGO-associated small RNAs are abundant and diverse. Here, we specifically probed the association of small RNAs in *Ascaris* PDE using a targeted and stage-, cellular-compartment-focused approach. We used antibodies to *Ascaris* Argonaute proteins and first performed immunohistochemistry (IHC) to identify Argonautes associated with mitotic chromosomes during PDE. We then carried out Argonaute-IP of the WAGOs on chromatin followed by dual RNA sequencing during PDE (DuoRIP-seq where both small RNAs and their target RNAs are sequenced). Overall, our data reveal distinct, region-specific WAGO-siRNA complexes during *Ascaris* PDE and demonstrate that small RNAs specifically associate with eliminated and retained chromatin.

## Results

### *Ascaris* WAGO-2 and WAGO-3 differentially localize to retained and eliminated DNA during PDE

We recently developed antibodies to seven highly expressed *Ascaris* Argonaute proteins, including AsALG-1 that binds to miRNAs, the testis-specific AsALG-4 associated with 26G-RNAs, and all five WAGOs (AsWAGO-1, AsWAGO-2, AsWAGO-3, AsCSR-1, and AsNRDE-3) that are associated with 22-24G-RNAs ^37^. Using these antibodies, we examined their localization within early *Ascaris* embryos during PDE. Because PDE occurs between the 4- and 16-cell stages and is restricted to pre-somatic lineages ^5^, we focused on 4-cell embryos, where 3 of 4 blastomeres undergo PDE. Immunohistochemistry staining revealed that all five WAGOs are present in both the cytoplasm and nucleus of *Ascaris* embryos (Fig. S1). AsCSR-1, AsWAGO-1, and AsNRDE-3 did not show any specific association with condensed chromosomes during PDE mitoses (Fig. S1). In contrast, the remaining two WAGOs are differentially and selectively enriched on condensed chromosomes during mitoses. Intriguingly, AsWAGO-2 is on retained, condensed chromosomes during PDE mitoses. Notably, it is not seen on condensed chromosomes during non-PDE mitoses (Fig. 1). In contrast, AsWAGO-3 is enriched on condensed and eliminated chromatin during PDE mitoses (Fig. 1). Even though only a small fraction (∼5%) of embryonic cells are in metaphase at any given time, our analysis of multiple embryos indicates these WAGO localization patterns occur across all mitotic cells during PDE (Fig. S1). Argonautes from the ALG clade (AsALG-1 and AsALG-4) did not stain condensed chromatin during PDE. Overall, two *Ascaris* WAGOs are specifically and differentially enriched on retained (AsWAGO-2) and eliminated (AsWAGO-3) DNA during PDE.

**Figure 1.**
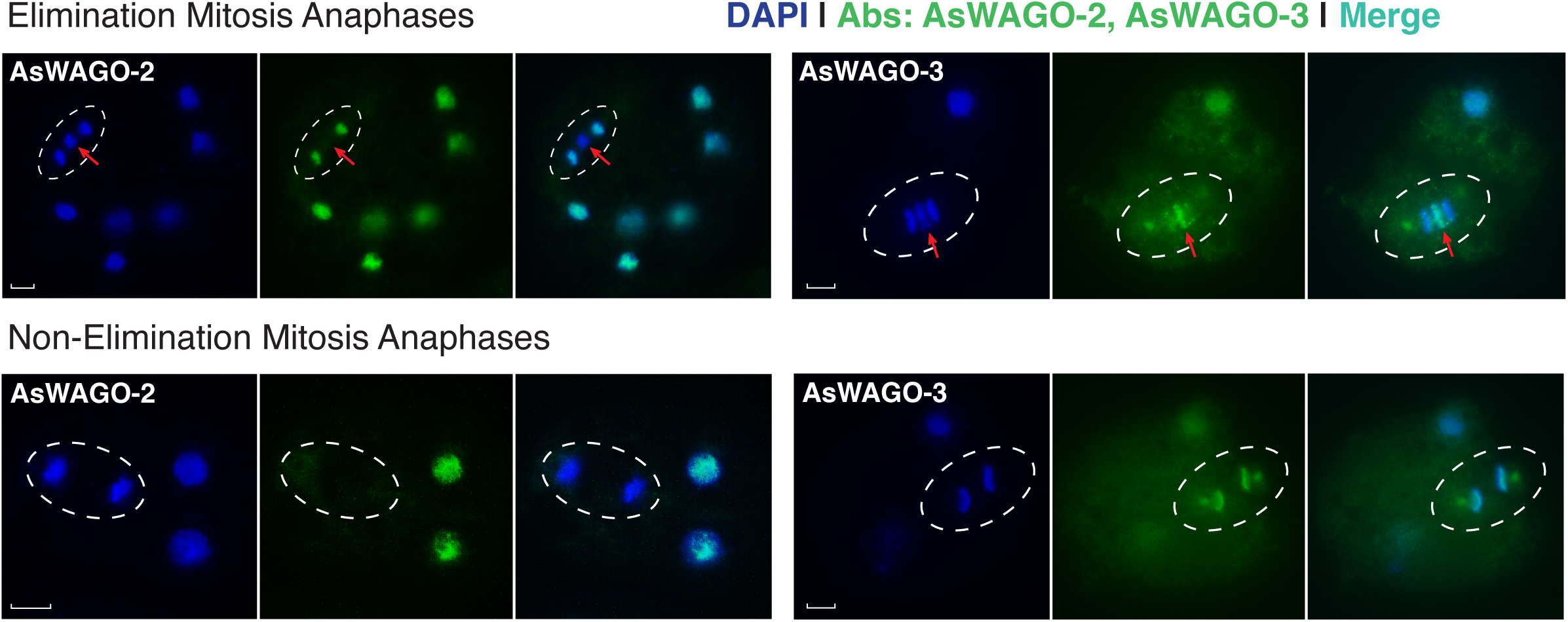
*Ascaris* WAGO-2 localizes to retained DNA during programmed DNA elimination mitoses, while WAGO-3 is enriched in eliminated DNA. Immunohistochemistry staining of early *Ascaris* embryos undergoing DNA elimination using antibodies against *Ascaris* Argonaute proteins. See Fig. S1 for additional staining images. The white dashed oval highlights a mitotic cell. The red arrow indicates DNA that is not segregated during mitosis, remains at the metaphase plate, and will be eliminated. Scale bars represent 10 μm.

Thes staining patterns contrast with our previous work using total small RNA sequencing ^36^ or whole cell immunoprecipitation of Argonaute-bound small RNAs ^37^, which did not identify an association between small RNAs and retained or eliminated sequences. However, the distinct IHC staining patterns of AsWAGO-2 and AsWAGO-3 (Fig. 1) suggest a specific association between these Argonautes and PDE. While the embryos used in previous and the current sequencing experiments were developmentally synchronous, only about 5% of the 4-cell embryos are in metaphase undergoing PDE, and one of the four blastomeres represents the germline that does not undergo DNA elimination. In addition, whole cell analyses include substantial contributions of cytoplasmic Argonautes and their small RNAs. Together, these factors likely reduced the signal-to-noise ratio, making it difficult to discern any potential association between small RNAs and PDE.

### Cellular fractionation and *Ascaris* WAGO immunoprecipitation

To further examine if WAGOs and their associated small RNAs localize to condensed chromosomes during PDE at sequence level, we performed subcellular fractionation to enrich for chromatin-associated AsWAGO-2 and AsWAGO-3 complexes, followed by sequencing (Fig. 2A). The standard RIP approaches recover the Argonaute-bound small RNAs but not their targets, thus they do not contain information about the enriched siRNA targets directly from the Argonaute-siRNA-target complex. We developed an RNase-assisted co-immunoprecipitation strategy ^39^ that exploits the fact that Argonaute proteins protect both their bound siRNAs and the directly paired region of their target RNAs from nuclease digestion (Fig. 2A). We termed this method DuoRIP-seq – a native, fractionation-based RIP adapted from the Fr-iCLIP protocol ^39,40^ (see Materials and Methods). DuoRIP-seq enables simultaneous recovery of (1) Argonaute-bound small RNAs (18-30 nt) and (2) the protected fragments of their target RNAs (sense, 18-150 nt) from the same immunoprecipitate, allowing direct, integrated analysis of their interactions (see Fig. 2A and Fig. S2) and enabling us to evaluate the levels and dynamics of these RNAs simultaneously. A 5’-independent library strategy was used to capture small RNAs with 5’-triphosphates or other non-canonical 5’-ends ^36,37^. RNA-seq on subcellular fractions confirmed strong enrichment for nuclear/chromatin RNAs, reflected by their higher intron/exon ratios (Fig. 2B-C). A comparison of Input and AsWAGO-3 IP (chosen for its strong target RNA association) also showed that DuoRIP-seq successfully captures both Argonaute-bound siRNAs and the corresponding chromatin-associated RNA fragments (Fig. 2B-C).

**Figure 2.**
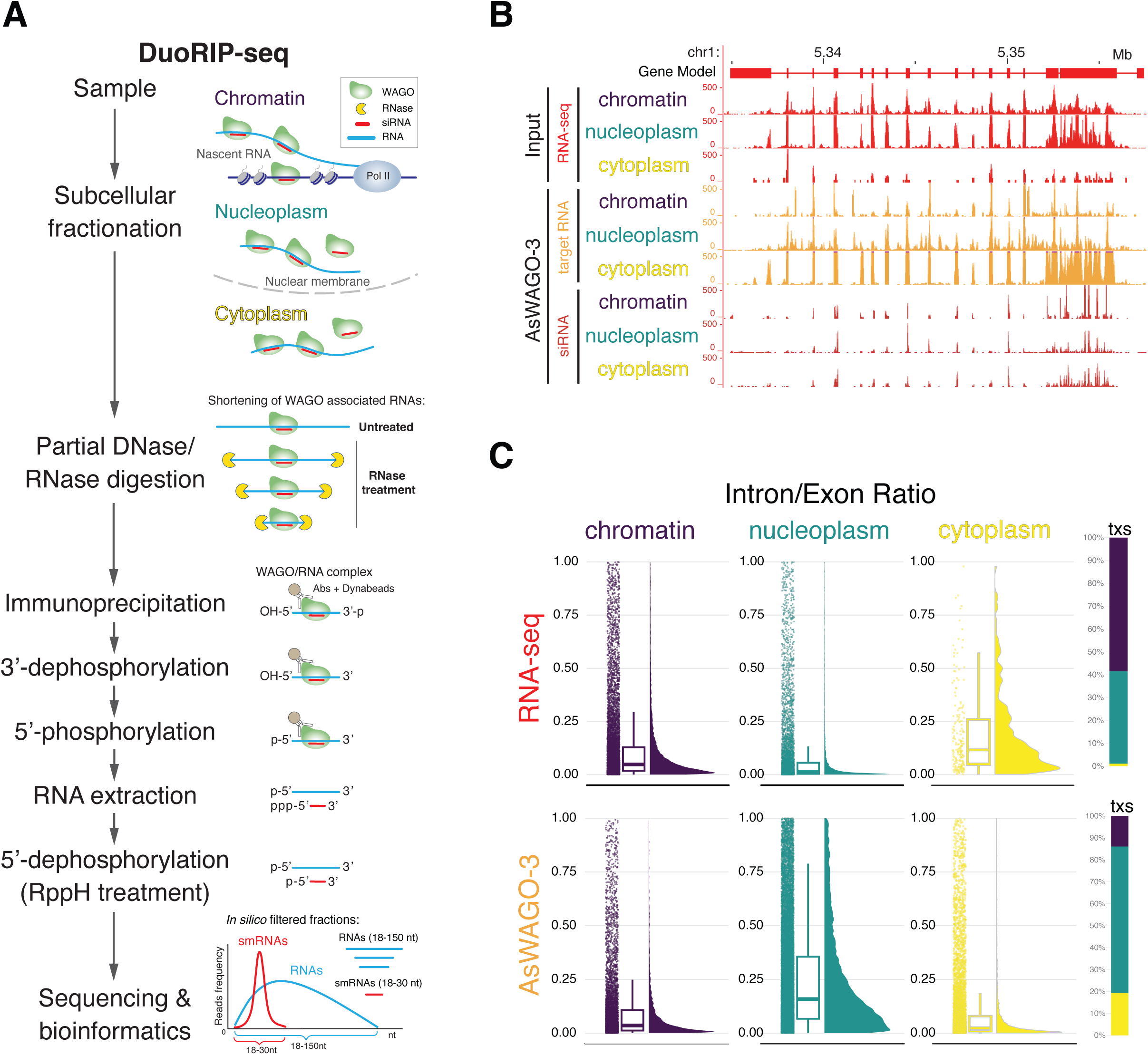
Enrichment of *Ascaris* Argonaute-associated RNAs in subcellular fractions. **A.** DuoRIP-seq overview. The key steps are illustrated. Chromatin, nucleoplasm, and cytoplasm were analyzed. Fragments of RNAs associated with Argonautes are protected from RNase digestion. Thus, the RNase capture assay identifies both small RNAs and their target RNAs associated with Argonautes. Protected small RNAs and their targets are distinguished by their size and strand orientation (see Fig. S2 for bioinformatic details on the discrimination of small RNAs and target RNA reads). **B.** Enrichment of target RNA reads in chromatin and nucleoplasm. Genome browser view of an exemplary gene showing total RNA from input (red), target RNA reads from AsWAGO-3 immunoprecipitation (orange), and small RNAs associated with AsWAGO-3 (dark red) from the three fractions. Note the enrichment of target RNA reads in nuclear fractions (intron reads) as well as enrichment of target RNA reads in AsWAGO-3 IP in the nucleoplasm **C.** Meta-analysis of target RNA read abundance using intron-to-exon (I/E) ratios in total RNA and AsWAGO-3 IP RNA. Raincloud plots display the I/E ratio distributions for individual transcripts across the three fractions. Higher I/E ratios, particularly in the nucleoplasm IPs, indicate enrichment of target RNA reads in association with AsWAGO-3. Vertical bars (right) represent the proportion of target RNA transcripts across subcellular fractions (in percentages). The intron and exon signals were normalized to their length and transcripts.

### *Ascaris* WAGO-2 and WAGO-3 bind to specific sets of small RNAs with distinct targets

We previously demonstrated the specificity of the *Ascaris* WAGO antibodies through Argonaute-IP and small RNA sequencing in germline tissues and embryos ^37^. Here, we used DuoRIP-seq to capture both Argonaute-bound small RNAs and their target RNA fragments in nuclear and cytoplasmic fractions (see Fig. 2). We used two close homologs of the AsWAGO-2/3 as negative controls for the DuoRIP-seq: AsWAGO-1 as the control for AsWAGO-2 and AsCSR-1 as the control for AsWAGO-3 because each pair is associated with similar classes of small RNAs and has similar target profiles ^37^, yet, the antibody staining showed that neither of these two control WAGOs is associated with condensed chromosomes during PDE (Fig. S1). Analysis of *Ascaris* WAGO-associated small RNAs from the different subcellular fractions shows that overall, the length distribution of small RNAs and their target RNAs for each AsWAGO (Fig. 3) is distinct and consistent with previous RIP-seq results using whole cells ^37^. In general, AsWAGO-3 and AsCSR-1 are associated with 23G- and 24G-RNAs and share many mRNA targets. Interestingly, a subset (∼25%) of AsWAGO-3 also binds to another type of siRNAs that targets primarily repeats in chromatin, while its counterpart AsCSR-1 almost exclusively (∼90%) targets mRNAs. For AsWAGO-1 and AsWAGO-2, they mainly bind to 22G-RNAs and share targets of repeats in genomic regions enriched for WAGO-associated siRNAs ^37^. Overall, our data illustrate distinct WAGO-associated siRNAs enriched in different subcellular fractions during *Ascaris* PDE.

**Figure 3.**
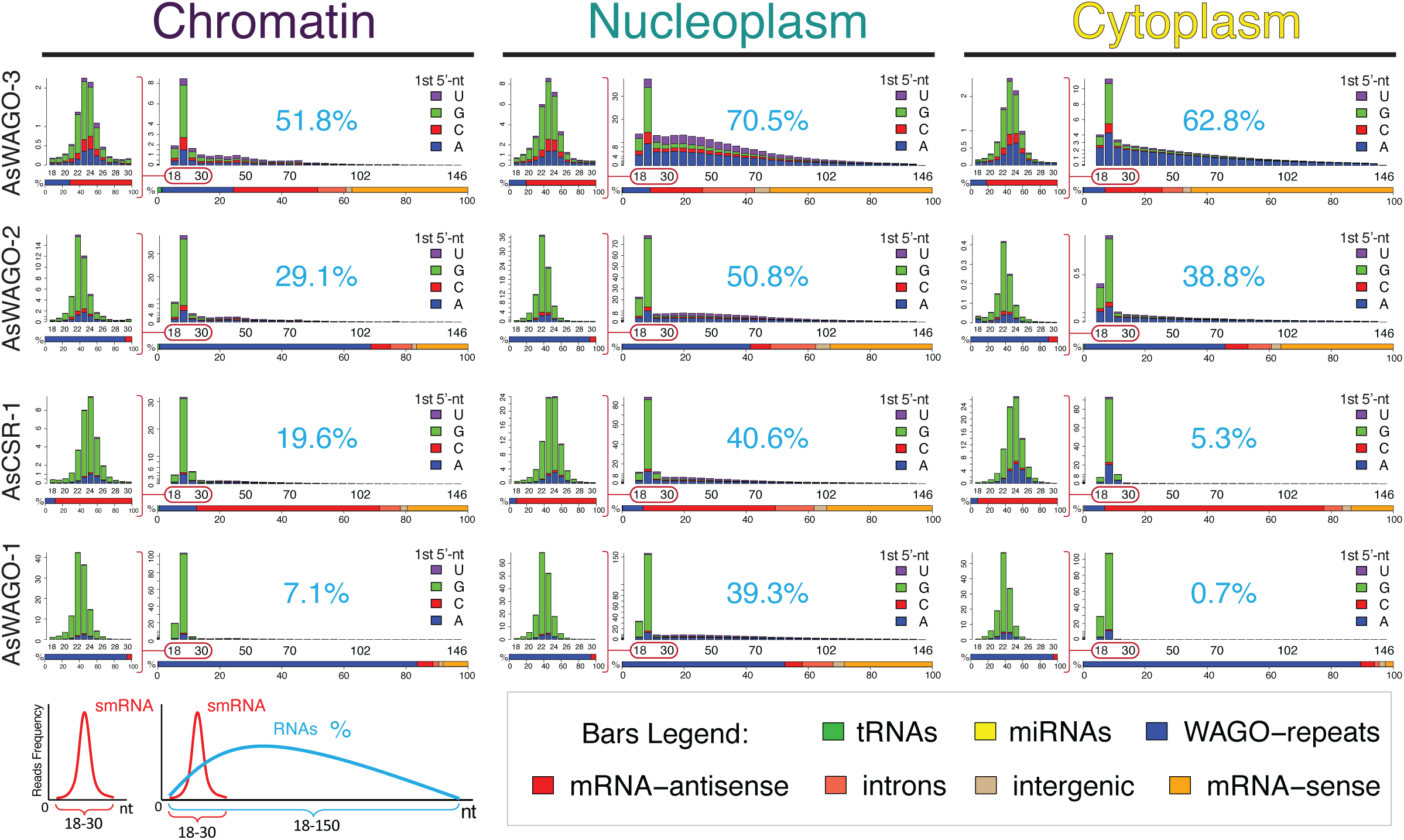
*Ascaris* Argonautes are associated with specific small RNAs and their targets. RNAs from four Argonaute antibody immunoprecipitations were sequenced, mapped, and categorized in chromatin, nucleoplasm, and cytoplasmic subcellular fractions (see methods). For each IP experiment, the size distribution on the left illustrates the frequency of small RNAs from 18 to 30 nt. The plot on the right shows the frequency of all WAGO-associated RNAs - siRNAs and their potential target RNAs - at a 4-nt window length (from 18 to 150 nt). Red ovals indicate the siRNAs. The percentage shown in blue represents the fraction of target-RNA reads relative to the total reads (siRNA + target RNAs) in each sample. RNAs starting with A, C, G, and U are color-coded. Two or more biological replicates that show consistent profiles were combined, except for the cytoplasmic samples. Below each size distribution plot are bars representing the siRNAs, their targets and their overall percentage of the small RNAs reads, including (1) tRNAs; (2) miRNAs; (3) siRNAs to WAGO-repeats (repetitive sequences and mobile elements targeted by AsWAGO-1, AsWAGO-2, and AsNRDE-3); (4) siRNAs antisense to mRNAs; (5) siRNAs matching introns; (6) siRNAs matching intergenic regions; and (7) sense small RNAs corresponding to mRNAs. Bars represent RNA categories for both siRNA (18-30nt) and RNA fragments (18-150 nt). See Fig. S4 for additional data on the size distributions and targets from biological replicates.

Our DuoRIP-seq also captures RNA fragments associated with the Argonautes (sense RNAs; Fig. 2A). These RNAs with lengths of 18 to 150 nt (Fig. 2A) are hereafter called target RNAs. Due to the continuous RNase digestion during the experimental procedure, the length distribution of the target RNAs varies among these libraries (Fig. 3). In addition, the percentage of reads for target RNAs relative to small RNAs differ drastically across the libraries, ranging from ∼0.7% to 70% (Fig. 3). This difference of target RNAs from DuoRIP-seq reflects the dynamic nature of RNAs that these WAGOs target across these cellular fractions. Notably, AsWAGO-2 and AsWAGO-3 have a higher percentage of target RNAs in chromatin compared to AsCSR-1 and AsWAGO-1, suggesting that AsWAGO-2 and AsWAGO-3 are highly enriched on chromatin and may target chromatin-bound RNAs. Interestingly, all WAGOs have more target RNAs in the nucleus than in the cytoplasm (Fig. 3), a reflection that the Ago-siRNA complexes could interact more in general with RNA transcripts within the nucleus. Overall, our DuoRIP-seq data identified specific AsWAGO associated siRNAs and their target RNAs in subcellular fractions.

### AsWAGO-3 associated siRNAs and their target RNAs are enriched in the eliminated DNA

To determine if AsWAGO-3 and its associated RNAs are enriched in chromosomal regions that will be eliminated, as demonstrated by IHC (Fig. 1), we mapped these RNAs to *Ascaris* retained and eliminated genomic sequences ^23^ and determined their target genes and repeats (Figs. 4A, 4C, and 4E). For the eliminated regions, both AsWAGO-3-associated siRNA and their target RNAs were predominantly enriched in the nuclear fractions, particularly in the nucleoplasm (Figs. 4B and 4D). In contrast, for the retained regions, AsWAGO-3 siRNAs and target RNAs showed no enrichment in the nuclear fractions (Figs. 4B and 4D). AsWAGO-3-associated siRNAs and their targeted RNAs are highly enriched in the eliminated regions within the nucleus, particularly for transcripts with introns exhibiting intron retention (Fig. 2C; compare AsWAGO-3 and input nucleoplasm).

**Figure 4.**
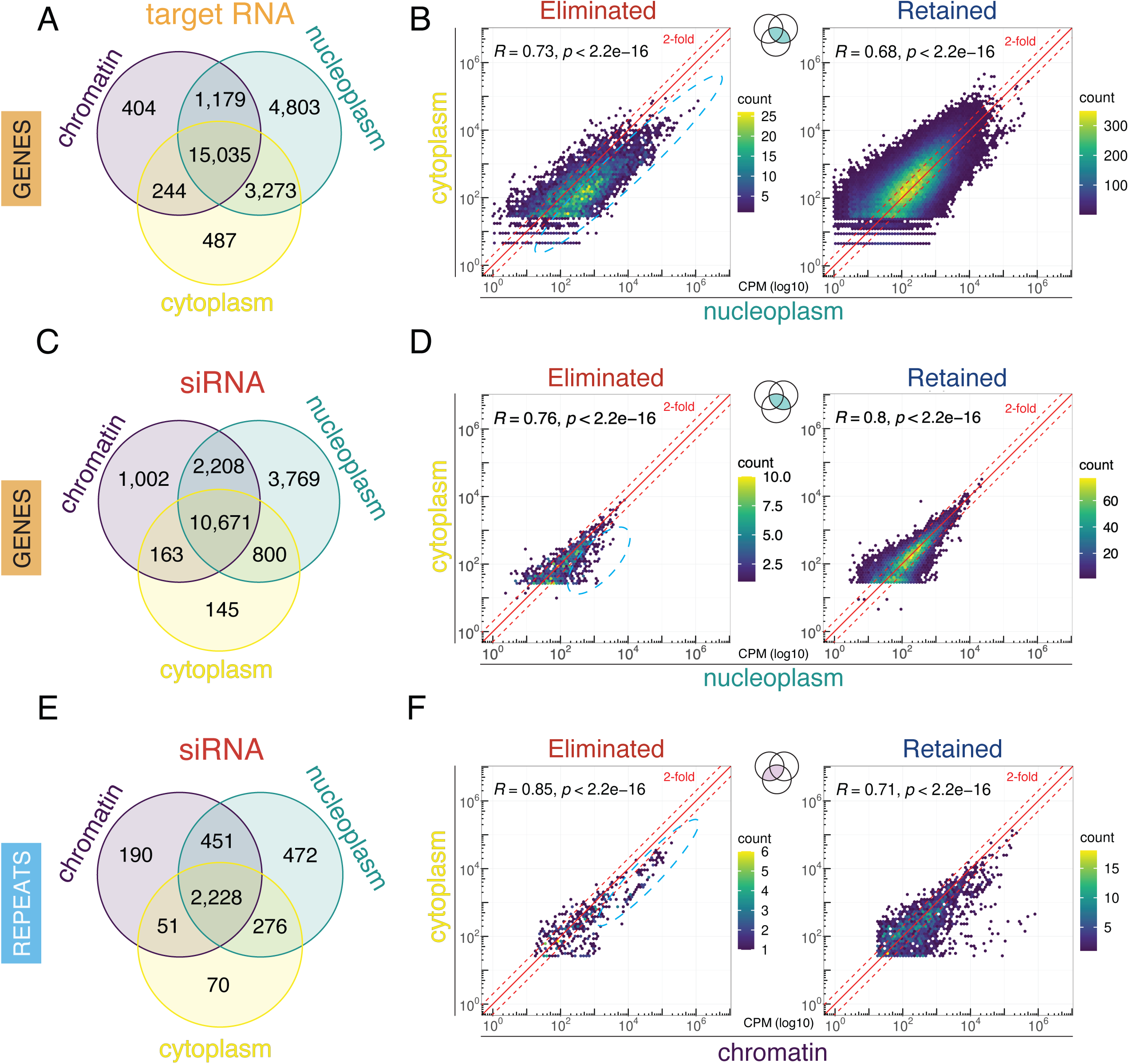
AsWAGO-3 associated siRNAs and associated RNAs are enriched for eliminated DNA. **A, C, E.** Numbers of genes (A, C) and repeats (E) targeted by AsWAGO-3 associated RNAs in the chromatin, nucleoplasm, and cytoplasm fractions. Targets were defined by ≥2-fold enrichment relative to input (see Methods). **B, D.** Scatterplots comparing RNA levels in the nucleoplasm and cytoplasm for each gene. Enrichment of AsWAGO-3–associated target RNA (B) and siRNA (D) in the nucleoplasm is especially pronounced for eliminated genes, which are highlighted with dashed line ovals. siRNAs (D) are shown for >2-fold enriched RNA targets in eliminated and retained in (B); for unfiltered siRNAs, see Fig. S5. **F.** Comparison of repeat-associated siRNAs in chromatin vs. cytoplasm. The data are in log scale of count per million reads, derived from early embryos (4-6 cell; 65hr) undergoing PDE. Point density is indicated by gradient color bars in scatterplots (B, D, F). Note the scales differ across plots. The siRNAs and target RNAs enriched in the nuclear fraction and targeting eliminated regions are highlighted with a dashed line oval. Color-coded Venn diagram icons indicate the subset of data represented in each panel. See Figure S5 & S8B for the complete AsWAGO-3 associated siRNA and target RNA datasets.

Although nuclear AsWAGO-3-associated siRNAs primarily target coding sequences (Figs. 3, 4A, and 4C), we identified repeats in the eliminated genomic regions that are also targeted by a subset of highly expressed siRNAs in the chromatin fraction (Figs. 4E and 4F). Unlike the highly transcribed genes that are targeted by relatively few siRNAs, these repeats exhibit low RNA expression, yet are targeted by highly abundant siRNAs (Fig. 4F, dashed line oval), suggesting AsWAGO-3’s specificity in targeting these repeats. About half of all *Ascaris* eliminated DNA consists of a 121-bp repeat ^21–23^. However, consistent with our previous total small RNA sequencing data ^36,37^, AsWAGO-3-associated siRNAs did not target the 121-bp repeat. Overall, the data indicate that AsWAGO-3-associated siRNAs are more abundant in the eliminated than retained genomic regions during PDE.

### AsWAGO-3 targeted eliminated repeats are mainly located in the middle of the chromosomes

To further examine the highly expressed AsWAGO-3 siRNAs targeting repeats in the eliminated regions (Fig. 4F), we first determined the chromosomal locations of these repeats. In *Ascaris*, most repetitive sequences reside within eliminated genomic regions and are often enriched in subtelomeric regions ^23^. However, some internal eliminated genomic regions are also enriched with repeats. Our comparative genomic analysis of several ascarid species suggested that internally eliminated DNA is derived from the end fusion of ancestral chromosomes to form their current germline chromosomes (Fig. 5A), and they are often associated with H3K9me3 ^25^. Remarkably, we found that the AsWAGO-3 targeted repeats are predominantly localized within these internally eliminated regions (Fig. 5B, track 1). Meta-analysis of the AsWAGO-3 chromatin fraction confirms internal eliminated regions are enriched with siRNAs that target these repeats (Fig. 5C). In contrast, AsWAGO-3 associated siRNAs that target genes are primarily present in terminal eliminated genomic regions (Fig. 5C). This targeting of the repeats in the middle of the chromosomes may be an adaptation of AsWAGO-3 to suppress previous heterochromatic sequences at the ends of the chromosomes that are now in an euchromatic environment in the middle of the chromosomes (see Discussion).

**Figure 5.**
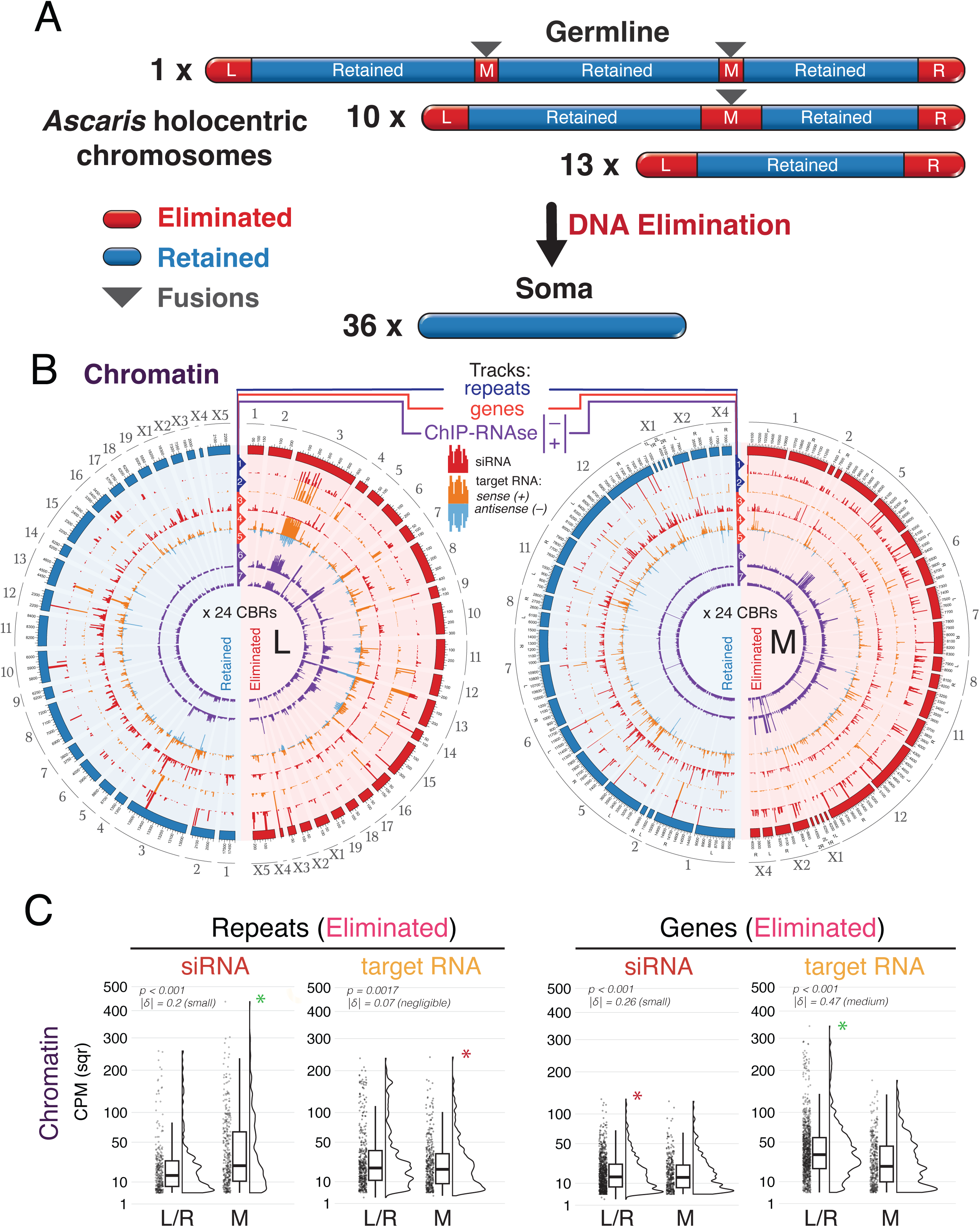
AsWAGO-3 associated small RNAs predominantly target DNA repeats in the middle of the chromosomes that are eliminated by PDE. **A.** Schematic representation of germline and somatic chromosomes illustrating terminal repeats (L and R) and those in the middle of chromosomes (M). Retained regions are shown in blue, and eliminated regions are shown in red. The numbers of chromosomes with and without eliminated DNA in the middle are indicated. The number of chromosomes increases from 24 in the germline to 36 in somatic cells following PDE ^23^. Presumptive ancestral chromosome fusion sites are marked with triangles [adapted from ^25^]. **B.** AsWAGO-3 associated small RNAs and target RNAs from chromatin fraction mapped to the left (L) and the middle (M) eliminated regions. The right (R) eliminated region has a similar pattern to the left one (see Figure S6). The outer circle of the circos plots displays eliminated regions (red) and randomly selected retained regions (blue) with matching size from the same chromosomes (see Methods). From the outside to the inside of the circos plot are the following seven tracks: (1) siRNAs targeting repeats, (2) target RNAs matching repeats, (3) siRNAs targeting genes, (4) sense RNAs matching genes, (5) antisense RNAs matching genes, (6) AsWAGO-3 ChIP-seq, and (7) AsWAGO-3 ChIP-seq with RNase treatment. The tracks are normalized as count per million reads (CPM). **C.** Levels of AsWAGO-3-associated siRNAs and target RNAs (CPM, square root scale) in eliminated regions are shown in the raincloud plots. siRNAs targeting repeats are significantly higher in the middle (M, green *) than at the ends (L/R) of the chromosomes. The eliminated regions in the middle of chromosomes also show a higher level of target RNAs (red *). Conversely, siRNAs targeting genes are more enriched at the ends (L/R, red *) than in the middle (M) of the chromosomes. Regions eliminated at the ends of chromosomes show much higher levels of target RNAs (L/R, green *). Green * reflects more abundant reads compared to red *. The patterns for the nucleoplasm fraction are similar (see Fig. S7). See also Figure S7 for comparison with nucleoplasm, cytoplasm fractions. The data are shown in a square root scale of CPM. Wilcoxon test (two-tailed) was used for significance; effect sizes are given as Cliff’s delta.

### AsWAGO-3 binds to chromatin through RNA-mediated interactions

Our data established an association between AsWAGO-3, chromatin, and eliminated DNA (Fig. 1 and Fig. 4). To further validate the chromatin association, we carried out ChIP-seq analysis using AsWAGO-3 antibodies. Our results showed an average of ∼1.6X enrichment of AsWAGO-3 on eliminated DNA compared to retained DNA (Fig. 6A). In addition, the identified ChIP-seq peaks from both internal and terminal eliminated regions are significantly broader than the retained regions (Fig. 6B). Consistently, IHC of the histone mark H3S10P, which stains the eliminated DNA during PDE mitoses ^23^, has a similar ChIP-seq profile to AsWAGO-3 (Fig. 6A). Overall, these data support our IHC staining and DuoRIP-seq data (Fig. 1 and Fig. 4), indicating that AsWAGO-3 is associated with chromatin and is enriched on eliminated DNA.

**Figure 6.**
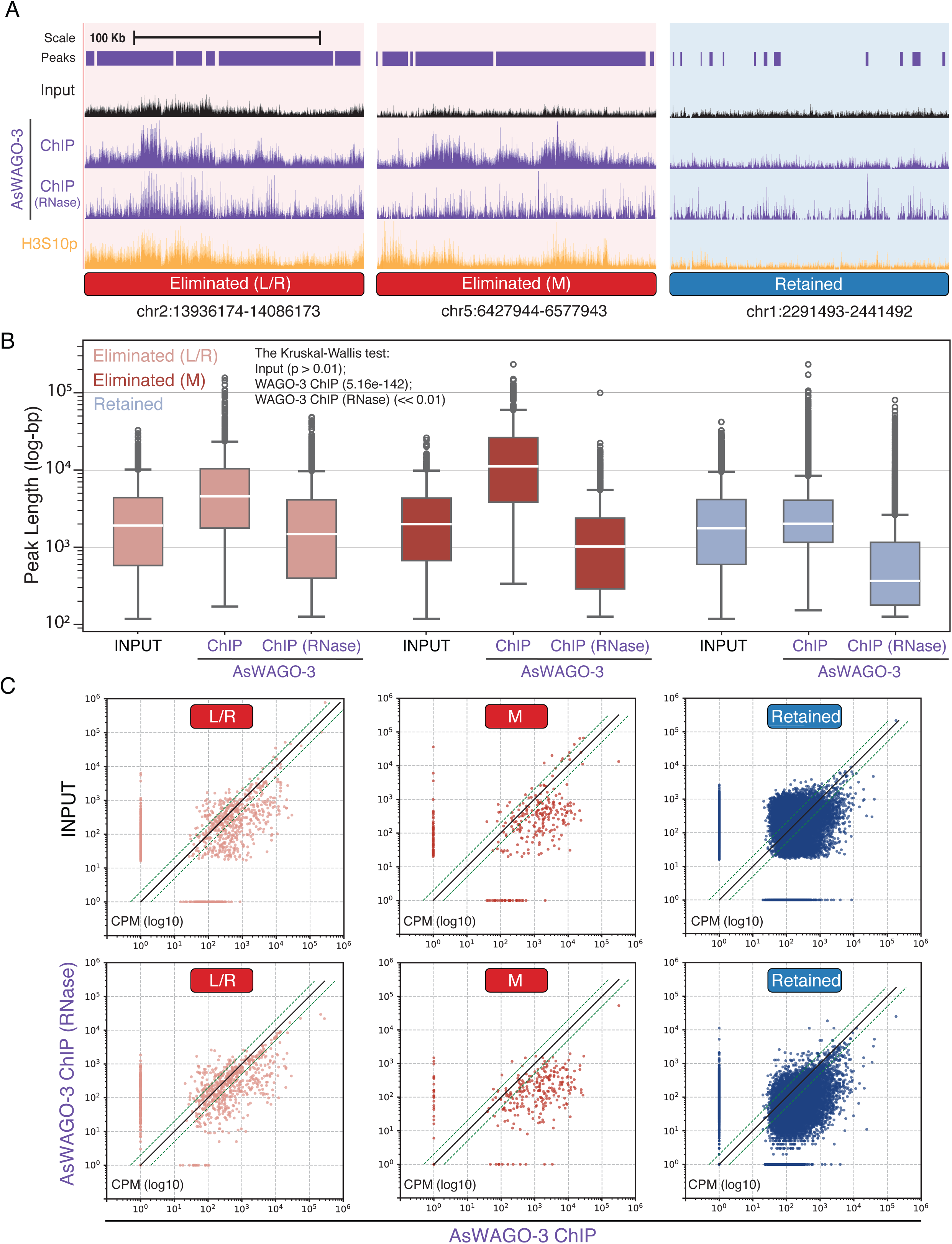
AsWAGO-3 binds to chromatin through RNA-mediated interactions and is enriched for eliminated regions. **A.** Representative regions of the *Ascaris* genome containing both eliminated and retained DNA, with chromatin Input (top), AsWAGO-3 ChIP-seq with and without RNase treatment (middle), and H3S10p ChIP-seq (bottom) from 4–6 cell embryos (65 h). L/R and M indicate terminal and internal eliminated regions of chromosomes. **B.** Boxplots showing the length distributions of AsWAGO-3 ChIP-seq peaks (y axis in log value) in *Ascaris* early embryos undergoing PDE. The peaks were called with MACS3 ^69^ (see methods). Data are shown for genomic regions that are eliminated from the chromosome arms (L/R – light red), eliminated from the internal region (M – dark red), and retained (blue). The Kruskal–Wallis test was used to assess whether peak length distributions significantly differ. **C.** AsWAGO-3 association with the eliminated chromatin is dependent on RNA. Input chromatin or RNase-treated AsWAGO-3 ChIP-seq data (y-axis) were compared to AsWAGO-3 ChIP-seq data (x-axis). The data were normalized as counts per million (CPM) and were binned into 1 kb windows across genomes. RNase-treated data were from the combined three biological replicates.

To further assess if AsWAGO-3 interacts with the chromatin through its siRNA binding to nascent RNAs, we treated the samples with RNase, followed by ChIP-seq. RNase treatment significantly reduced AsWAGO-3 on the chromatin across the genome (Fig. 6A-C). The level of reduction varies in different parts of the genome, with the internal eliminated regions (M) showing the most significant decrease of AsWAGO-3 after RNase treatment, followed by the terminal eliminated regions (L/R) and the retained regions (Fig. 6B-C). Overall, the ChIP-seq results confirm that AsWAGO-3 is associated with chromatin, and this association requires RNA-mediated interactions, likely through the complementary binding of antisense small RNA to nascent RNAs.

### AsWAGO-2 siRNAs and their association with the retained DNA

Most AsWAGO-2 siRNAs target repetitive elements, which are primarily found in genomic regions eliminated during PDE. Consequently, immunoprecipitation of AsWAGO-2 from whole-cell lysates without subcellular fractionation recovered siRNAs predominantly mapping to eliminated regions (Fig. 3). However, immunostaining revealed that AsWAGO-2 localizes specifically to condensed, retained chromosomes during PDE mitoses. Intriguingly, AsWAGO-2 also localizes to interphase nuclei, but not to chromosomes in non-PDE mitoses (Fig. 1 and Fig. S1). This apparent discrepancy between AsWAGO-2 cellular localization (mostly visible in condensed retained chromosomes) and its associated siRNA targets in the genome (mostly in eliminated regions) suggested that a subset of AsWAGO-2 complexes might transiently interact with retained chromatin, and these complexes would likely be underrepresented in total cellular extracts.

To examine this, we analyzed subcellular fractions using DuoRIP-seq. While AsWAGO-2-associated siRNAs and their target RNAs were overall highly concentrated in the chromatin and nucleoplasm fractions (Fig. 7A), within the chromatin fraction, we identified a distinct group of AsWAGO-2-associated siRNAs and target RNAs that were specifically enriched in retained DNA regions (Fig. 7B). Notably, these chromatin-enriched siRNAs largely overlapped with target RNAs derived from the same repetitive loci (Fig. 7B, green dots in the retained regions), suggesting that both these AsWAGO-2 siRNAs and their targeted RNAs are enriched in the retained chromatin fractions, likely in the form of Argonaute-siRNA-target complexes engaging with chromatin. These findings reconcile the immunohistochemistry data, indicating that while most AsWAGO-2-associated siRNAs target repeats in eliminated DNA, a specific subset marks repeats within retained chromatin during PDE. This subset may represent a transient but functionally important interaction of AsWAGO-2 with retained chromatin through its binding of repeat-derived RNA transcripts.

**Figure 7.**
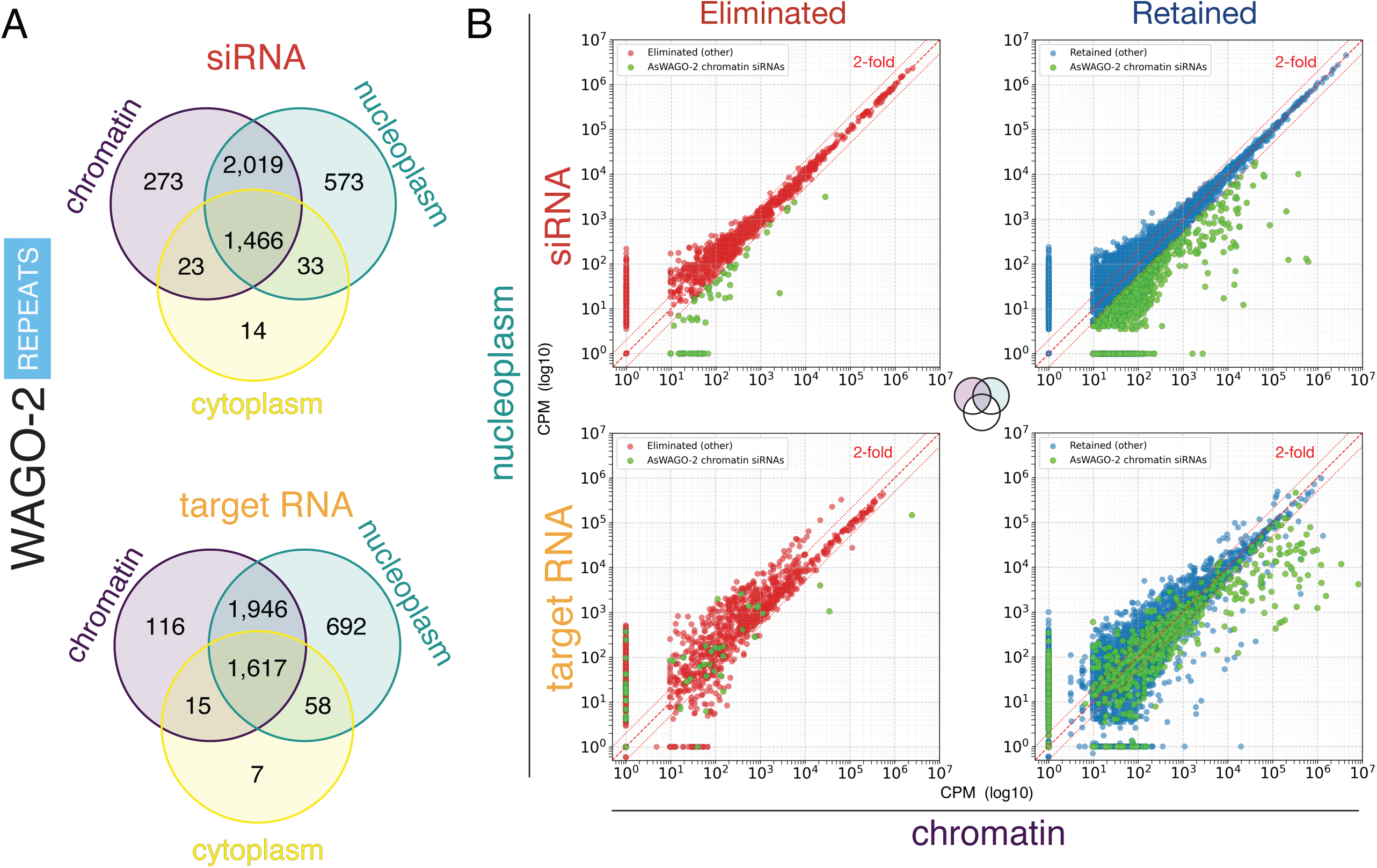
AsWAGO-2–associated siRNAs and associated RNAs are enriched for retained genome regions. **A.** Venn diagrams showing unique and overlapping AsWAGO-2–associated siRNAs (top) and target RNAs (bottom) matching repeats in the chromatin, nucleoplasm, and cytoplasm fractions. **B.** Enrichment of siRNAs and target RNAs in the chromatin over nucleoplasm fractions for the retained genomic regions. The abundance of AsWAGO-2–associated siRNAs (top) and RNAs (bottom) mapped to eliminated (left) and retained (right) genomic regions (in CPM, log₁₀ scale). Red and blue points represent eliminated and retained loci, respectively; green marks siRNAs or RNAs associated with chromatin-enriched AsWAGO-2 siRNAs targets that are >2-fold enriched in chromatin. See Figure S8A for the complete datasets for AsWAGO-2 associated siRNA and target RNAs.

## Discussion

Small RNAs and their effector Argonaute proteins play diverse regulatory roles in gene expression, genome defense, and epigenetic control. Argonautes are generally classified into two major clades: AGO and PIWI ^41,42^. In nematodes, however, an additional clade of expanded worm-specific Argonaute (WAGOs) is highly abundant. These WAGOs regulate gene expression through gene licensing, fine-tuning, and silencing mechanisms ^43–47^. Despite their importance in *C. elegans*, relatively little is known about WAGO function in other nematodes ^37,48^.

Due to the high abundance and complexity of small RNAs in nematodes, our earlier sequencing study examining small RNAs before, during, and after PDE failed to establish a specific association between small RNAs and *Ascaris* PDE ^36^. In this study, we used several approaches to increase signal-to-noise in our analyses. First, we generated antibodies to individual WAGOs to define specific small RNA populations associated with individual Argonautes. Second, we applied cell fractionation to enrich for nuclear associated WAGOs and their small RNAs. Third, we sampled the WAGO-protected RNAs to simultaneously identify siRNAs and their targets. Finally, we employed multiple methods, including IHC, DuoRIP-seq, and ChIP-seq, to corroborate our results. These improvements enabled us to enrich for small RNAs associated with PDE.

We identified two *Ascaris* WAGOs, AsWAGO-2 and AsWAGO-3, that are associated with PDE. During PDE mitoses, AsWAGO-2 was found by IHC to be enriched on chromosomes that are retained, while AsWAGO-3 was primarily associated with eliminated chromosome regions. Notably, AsWAGO-3-associated siRNAs predominantly targeted repeats located within internally eliminated regions that arise from ancestral chromosome-end fusions. Despite AsWAGO-2’s predominant association with repeats in eliminated regions, a subset of AsWAGO-2-associated siRNA complexes transiently engages retained chromatin during PDE. These findings demonstrate that specific *Ascaris* WAGOs may contribute to PDE by reinforcing transcriptional repression of genes and repeats as well as responding to karyotype changes caused by ancient chromosome fusions.

### Enrichment of WAGOs and their associated small RNAs on retained and eliminated DNA

Our initial hypotheses for a potential role of Argonautes and small RNAs in *Ascaris* PDE were that small RNAs might target and enable sites for chromosomal breaks ^22^, depletion of CENP-A in regions of chromosomes that are eliminated ^19^, or targeting the major 121-bp tandem repeat that is eliminated ^23^. Although we used a 5’-independent library strategy to capture siRNAs with 5’ polyphosphates, certain siRNAs might be missed due to other modifications at their 5’, 3’, or internal regions. We also did not explore the possibility that mismatch tolerance for small RNA targeting as observed in *C. elegans* piRNAs ^49,50^. Nevertheless, our current data and analysis do not support that *Ascaris* small RNAs target chromosomal breakage regions, CENP-A-depleted regions, or the 121-bp repeat.

Our study shows that AsWAGO-3 is associated with eliminated chromosome regions and targets both genes and repeats. This association is supported by antibody staining (Fig. 1), DuoRIP-seq (Fig. 4 and Fig. 5), and ChIP-seq (Fig. 6). The expression of the AsWAGO-3 gene increases before and during PDE (Fig. S3), and the IHC staining data illustrate dynamic, region-specific chromatin localization (Fig. 1 and Fig. S1). Overall, these data support a link between AsWAGO-3 and PDE. AsWAGO-3 is selectively enriched in other specific regions of the eliminated chromosomes and may serve to silence certain genes that reside at the chromosome ends, as well as specific repeats at the internal regions of chromosomes caused by ancient fusion events^25^.

Our previous IHC staining showed that H3S10P, a histone mark associated with condensed chromosomes, persists on eliminated DNA during mitoses ^23^. Immunogold labelling of H3S10P indicated that the eliminated DNA is engulfed into micronuclei. The persistence of the H3S10P on the eliminated chromatin is likely due to the failure or incomplete dephosphorylation of the mark within micronuclei ^23^. Since AsWAGO-3 and H3S10P have similar IHC staining patterns and ChIP-seq profiles (Fig. 6A), we speculate that the enrichment of AsWAGO-3 on the eliminated DNA could also be associated with H3S10P histone or other unidentified specific histone marks. Although the IHC showed AsWAGO-3 is on the eliminated DNA during PDE, we were not able to determine from either IHC or ChIP-seq if AsWAGO-3 is also enriched in the eliminated DNA before PDE mitoses.

Our analyses reveal that AsWAGO-2, primarily associated with repetitive sequences in eliminated regions, also interacts with a distinct subset of siRNAs and target RNAs originating from retained chromatin during PDE. Subcellular fractionation resolves the apparent discrepancy between sequencing and staining data, suggesting that AsWAGO-2 forms complexes not only in the nucleoplasm but also on chromatin. The enrichment of AsWAGO-2-associated siRNAs and their target RNAs in the retained DNA indicates that a subset of AsWAGO-2 complexes transiently interacts with active chromatin during PDE mitoses. These findings support a model in which AsWAGO-2 participates in multiple regulatory pathways of genome surveillance: the majority of its siRNAs may be used to silence the repeats in eliminated DNA, while a small fraction could be used to transiently engage with the repeats in the retained chromatin during PDE.

Although AsWAGO-2 and AsWAGO-3 are differentially enriched in retained and eliminated DNA during PDE, respectively, their expression and targeting is not restricted to PDE (Figs. 4, 5, and 7 and Fig. S4). AsWAGO-3 is expressed in the male germline and somatic tissues (intestine and carcass), and AsWAGO-2 is expressed in the female germline (Fig. S3) ^37^. This indicates that their primary roles may extend beyond PDE. Interestingly, the majority of their small RNA targets in diverse tissues overlap with those of other WAGOs, particularly AsWAGO-1 and AsCSR-1 (Fig. 3), suggesting these WAGOs may be functionally redundant. Moreover, AsWAGO-3 is expressed at levels 10-50 times lower than AsCSR-1, and AsWAGO-2 is expressed at levels 2-4 times lower than AsWAGO-1 (34), suggesting that AsWAGO-2 and AsWAGO-3 may supplement the functions of AsWAGO-1 and AsCSR-1. Furthermore, it is possible that AsWAGO-2 and AsWAGO-3 could be co-opted to play other roles in PDE.

Overall, we propose that siRNAs associated with AsWAGO-2 and AsWAGO-3 constitute an adaptive mechanism that acts in concert during PDE to transiently silence genes and repeats in both eliminated and retained chromatin. AsWAGO-3-associated siRNAs target two major classes of loci within eliminated regions: genes at the chromosome ends and repeats in the middle of the chromosomes. In contrast, AsWAGO-2-associated siRNAs transiently target a subset of highly expressed repeats in retained chromatin. These siRNA pathways may complement the function of PDE silencing to fine-tune gene and repeat repression without directly contributing to determining which sequences are retained or eliminated.

### Mobilization of AsWAGO-3 to silence ancestral repeats in the middle of *Ascaris* chromosomes

AsWAGO-3 associated siRNAs are enriched for the eliminated DNA. While the majority of these are low-frequency siRNAs that target genes, a subset are high-frequency siRNAs that target repetitive sequences (Fig. 4). Remarkably, these targeted repeats are predominantly located in the middle but not the ends of the chromosomes. This raises an intriguing question: why does AsWAGO-3 selectively target the rare internal repeats instead of the highly abundant terminal repeats? Our recent study comparing the genomes of three ascarids (*Ascaris, Parascaris,* and *Baylisascaris*) suggests that their germline chromosomes are the result of the fusion of smaller ancestral chromosomes ^25^. Our analyses indicate that the eliminated regions in the middle of the germline chromosomes originated from the ends of ancestral chromosomes. Notably, a conserved characteristic of nematode PDE is chromosome end-remodeling, where the ends of all chromosomes are removed during DNA elimination ^5^. Chromosome fusion internalizes the ancestral ends, creating genomic regions that may require additional regulatory control. Small RNA targeting of these internalized regions likely represents an adaptive mechanism to regulate sequences that were previously terminal but are now located in euchromatic and potentially more transcriptionally active genomic regions.

Typically, nematode chromosomes have more housekeeping genes in the middle and more repeats at their ends ^51^. These terminal repeats are often silenced by heterochromatin ^52^, likely due to telomere anchoring at the nuclear periphery ^53^. Following fusion, remnants of these ancient terminal repeats are incorporated into the middle of the chromosomes, which are likely more euchromatic. We propose that AsWAGO-3 has been mobilized to silence these repeats, thereby mitigating potential instability or inappropriate expression at fusion junctions. The nuclear localization of abundant AsWAGO-3 siRNAs and the low levels of target RNA expression support this model. It is also consistent with the specific targeting of the repeats in the middle but not the ends of the *Ascaris* germline chromosomes, where repeats are defined as WAGO-enriched loci identified in our previous study ^37^. Interestingly, small RNAs associated with AsWAGO-3 in the germline tissues are also enriched in these internalized repeats ^37^, consistent with targeted suppression of fusion-derived regions.

Chromosome fusion and fission events theoretically can be tolerated in nematodes due to the holocentric nature of their chromosomes ^54–56^. These events are a primary evolutionary force that shapes nematode genomes ^25,57–59^. However, these structural changes may introduce vulnerability, necessitating mechanisms to stabilize newly formed genomic regions. Worm-specific Argonautes, including AsWAGO-3, may offer a solution to these genome alterations. Nematode WAGO pathways are redundant, versatile, potent, and adaptable, capable of carrying out various functions ^33,35,60^. Consequently, they could serve as a convenient means to silence repetitive regions in the middle of the chromosomes. For instance, our previous study demonstrated that, in addition to AsWAGO-3, AsNRDE-3 can also alter its targets during gametogenesis, transitioning from primarily targeting repetitive regions in the mitotic zone to predominantly targeting genes in the meiotic zone ^37^. This shift of AsNRDE-3 targets in the germline may facilitate the clearance of mRNAs during late spermatogenesis. Interestingly, in *C. elegans*, the orthologous NRDE-3 is also observed to switch targets during embryogenesis, shifting from targeting germline-expressed genes during oogenesis to repressing recently duplicated genes and retrotransposons in somatic cells ^61^. Together, these observations highlight the adaptability of WAGO pathways in maintaining genome integrity. WAGO-associated siRNAs can not only license and fine-tune gene expression but also suppress potential deleterious effects arising from structural genome changes such as chromosome fusion.

### Small RNAs and *Ascaris* PDE

In ciliate PDE, PIWI Argonautes and their associated piRNAs scan the genome to mark retained or eliminated DNA ^14,27,28^. This scanning model inspired us to search for small RNAs in *Ascaris* that might play a similar role. Although *Ascaris* has lost the PIWI pathway, its diverse Argonautes and abundant siRNAs could have been adapted to contribute to PDE ^37^. While our data indicate a strong association between AsWAGO-3 and eliminated DNA, we also discovered that other genomic regions are targeted by AsWAGO-3, albeit at a lower level. Considering the adaptability of WAGOs and the unique characteristics of eliminated DNA, such as their association with histone marks (H3S10P) ^23^, 3D genome organization ^62^, and micronuclei ^23^, we propose that the observed association is more likely a consequence of AsWAGO-3’s response to the chromatin alterations in eliminated DNA, rather than a causal factor that marks these regions for elimination.

Similarly, for AsWAGO-2, we only observed a transient association with retained DNA during elimination mitoses. While our data do not directly demonstrate causality, the RNA-dependent chromatin association of AsWAGO-3, together with its enrichment on eliminated regions and targeting of specific transcripts, provides a mechanistic framework for how small RNAs could contribute to chromatin specification during PDE. Overall, we suggest that AsWAGO-2 and AsWAGO-3 contribute to nuclear silencing pathways activated in response to inappropriate expression, or to reinforce repeat silencing accompanying rather than determining DNA elimination.

## Conclusion

Nematodes have evolved a diverse class of Argonaute proteins known as WAGOs. These WAGOs serve a variety of regulatory functions and exhibit remarkable flexibility in their targets and modes of action. Here, we document that two *Ascaris* WAGOs are associated with PDE. Notably, AsWAGO-3 is strongly linked to repeats in the middle of chromosomes, targeting ancestral chromosome-end repeats that have become internalized through ancient fusion events. Together, AsWAGO-3 and AsWAGO-2 contribute to the silencing of specific loci in both eliminated and retained DNA, acting in concert with PDE. These findings highlight the plasticity of WAGO pathways, their contribution to gene and repeat silencing, and their potential roles in nematodes that undergo PDE.

## Materials and Methods

### Ascaris

*Ascaris* embryonation was conducted as previously described ^36,63^.

### Antibodies and immunohistochemistry

Polyclonal antibodies to *Ascaris* proteins AsALG-1, AsALG-4, AsCSR-1, AsWAGO-1, AsWAGO-2, AsWAGO-3, and AsNRDE-3 were generated and affinity-purified as previously described ^37^. *Ascaris* embryo immunohistochemistry was carried out as described ^63^, using a modified freeze-crack method to permeabilize and fix embryos.

### DuoRIP-seq overview

An overview of Duo-RIP-seq is illustrated in the flowchart of Fig. 2A. Briefly, *Ascaris* 4-cell embryos were fractionated into chromatin, nucleoplasm and cytoplasm using a urea-based extraction method. Each fraction was treated with DNase and a limiting amount of RNase I to enrich for Argonaute-protected RNAs, followed by immunoprecipitation using polyclonal anti-AsWAGO antibodies bound to Protein A Dynabeads. After sequential RNA end repair with T4 PNK, proteinase K digestion, and TRIzol extraction, samples were treated with RppH to convert siRNAs’ 5′ polyphosphates to monophosphates. The recovered RNAs were used to construct libraries using the Lexogen Small RNA-Seq Kit, enabling simultaneous profiling of small RNAs and short target RNA fragments by Illumina sequencing (see detailed steps below).

### Chromatin, nucleoplasm and cytoplasm fractionation

Fresh *Ascaris* 4-cell embryos were decoated ^36,63^ and directly homogenized in a metal dounce at 4°C. The nuclei were prepared as described ^64^ with an increase to 0.5% Triton X-100 to improve the purity. Nuclei and cytoplasm were collected from the pellet and supernatant after centrifugation at 1,200 x g for 15 minutes at 4°C. Both nuclei and cytoplasm were flash frozen in liquid nitrogen and stored at -80°C. To separate chromatin and nucleoplasm for each sample, 400 μl of frozen nuclei were thawed on ice for ∼10 minutes and resuspended in 4 ml Lysis Buffer (50 mM Tris-HCl, pH 7.4, 150 mM NaCl, 2 mM MgCl_2_, 0.5% NP-40 (v/v), and 1X protease inhibitor cocktail from Roche). Nuclei were incubated on ice for another 10 minutes, then centrifuged at 950 x g for 10 minutes at 4°C, and the supernatant discarded. The nuclear pellet was resuspended in 400 µl Buffer 1 (50% glycerol (v/v), 20 mM Tris–HCl pH 7.9, 75 mM NaCl, 0.5 mM EDTA, 0.85 mM DTT), followed by the addition of 3.6 ml Buffer 2A (20 mM HEPES pH 7.6, 300 mM NaCl, 0.2 mM EDTA, 1 mM DTT, 7.5 mM MgCl_2_, 1 M urea, 1% NP-40, 200 U of NxGen RNase Inhibitor from Lucigen). The samples were vortexed for 10 seconds and incubated on ice for 10 minutes. Chromatin was sedimented at 15,000 × g for 5 minutes at 4°C. The supernatant was transferred into clean 15 ml tubes and stored on ice (nucleoplasm fraction). For the chromatin pellets, 400 μl of Buffer 1 was added and 3.6 ml of Buffer 2B (20 mM HEPES pH 7.6, 300 mM NaCl, 0.2 mM EDTA, 1 mM DTT, 7.5 mM MgCl_2_, 1 M urea, 1.5% NP-40 (v/v), 200 U of NxGen RNase Inhibitor from Lucigen). Samples were vortexed for 10 seconds and incubated on ice for 10 minutes. The chromatin was re-sedimented at 15,000 × g for 5 minutes at 4°C. The supernatant was discarded, and the pellets were washed twice with 1.2 ml of Buffer 2A. Finally, the chromatin was sedimented at 15,000 × g for 5 minutes at 4°C. The chromatin fraction was resuspended in 1 ml of Buffer 3 (50 mM Tris-HCl, pH 7.4, 100 mM NaCl, 0.1% SDS, 0.5% Sodium deoxycholate, 200 U of NxGen RNase Inhibitor from Lucigen).

### DNase/RNase partial digestion

DNA and RNA in the chromatin, nucleoplasm, and cytoplasm fractions were partially digested before immunoprecipitation to make target proteins more accessible and to fragment long RNAs for sequencing along with small RNAs. Eight μl of Turbo DNase (16 U) (Invitrogen, catalog #AM2238) was directly added into each fraction: chromatin (∼1 ml), nucleoplasm (∼4 ml), and cytoplasm (∼4 ml), and was incubated at 37°C for 15 minutes, shaking at 1,100 rpm. After incubation, 20 μl of RNase I (Invitrogen, catalog #AM2294) was diluted 1:50 (40 U; determined empirically) in sterile PBS. After 4-5 minutes of incubation at 37°C with 1,100 rpm shaking, all samples were immediately transferred to ice for 5 minutes. Samples were centrifuged for 10 minutes at 22,000 x g at 4 °C to clear the lysates. Supernatants were carefully collected into fresh nuclease-free tubes. Trizol LS (Invitrogen, catalog #10296010) was added to the input samples while other samples were further processed for immunoprecipitation.

### Argonaute immunoprecipitation

Bead preparation was done according to iCLIP ^39^. 300 μl of protein A Dynabeads (Invitrogen, catalog #10001D) per experiment (100 µl for chromatin, nucleoplasm, or cytoplasm) were washed twice with Lysis buffer 2 (without protease inhibitor) (50 mM Tris-HCl pH 7.4, 100 mM NaCl, 1% Igepal CA-630, 0.1% SDS, 0.5% Sodium deoxycholate, 200 U of NxGen RNase Inhibitor from Lucigen). This and all following bead washing steps were performed with 900 μl of their respective buffers. The beads were resuspended in 300 µl Lysis buffer 2 (50mM Tris-HCl, pH 7.4, 100mM NaCl, 1% Igepal CA-630, 0.1% SDS, 0.5% Sodium deoxycholate, 200 U of NxGen RNase Inhibitor from Lucigen) with 2–10 µg antibody, and the tubes were rotated for 60 minutes at room temperature. Unbound antibodies were removed from the beads with 3 washes with 1 ml Lysis buffer 2 (50 mM Tris–HCl pH 7.4, 100 mM NaCl, 1% Igepal CA-630, 0.1% SDS, 0.5% Sodium deoxycholate, 200 U of NxGen RNase Inhibitor from Lucigen). Argonaut IPs were carried out with rabbit antibodies to *Ascaris* WAGO’s bound to Protein A Dynabeads, and rabbit IgG (Millipore) was used as a control. Samples were mixed with the beads and rotated overnight at 4°C. The beads were then placed on a magnet, and the supernatant was discarded. Beads were washed twice with High-salt buffer (50 mM Tris–HCl, pH 7.4, 1 M NaCl, 1 mM EDTA, 1% Igepal CA-630, 0.1% SDS, 0.5% sodium deoxycholate) followed by two washes with PNK buffer (20 mM Tris–HCl, pH 7.4, 10 mM MgCl_2_, 0.2% Tween-20). Beads were left in 1 ml PNK buffer before proceeding to treat the RNA ends as described ^39^.

### RNA 3’-end dephosphorylation

Beads were placed on a magnetic rack, and the supernatant was discarded. Then beads were resuspended in 20 μl of the following mixture: 4 μl 5x PNK pH 6.5 buffer (350 mM Tris–HCl, pH 6.5, 50 mM MgCl_2_, 5 mM dithiothreitol), 0.5 μl T4 polynucleotide kinase (5 U) (PNK, NEB, catalog #M0201S), 0.5 μl NxGen RNase Inhibitor (20 U) from Lucigen, and 15 μl of nuclease-free water (Ambion). The reaction was incubated for 20 minutes at 37°C in a thermomixer at 1,100 rpm. After incubation beads were washed with 1 ml of PNK buffer (20 mM Tris–HCl, pH 7.4, 10 mM MgCl_2_, 0.2% Tween-20) then with 1 ml of High-salt buffer (50 mM Tris– HCl, pH 7.4, 1 M NaCl, 1 mM EDTA, 1% Igepal CA-630, 0.1% SDS, 0.5% sodium deoxycholate) and finally twice with 1 ml of PNK buffer.

### RNA 5’-end phosphorylation

Beads collected after 3’-end dephosphorylation were resuspended in 20 μl of PNK mix: 0.25 μl PNK (2.5 U), 1 μl 10 mM ATP, 2 μl 10x PNK buffer, 0.25 μl NxGen RNAse Inhibitor (Lucigen) (40 U/µl), and 16.5 μl of nuclease-free water (Ambion). The reaction was incubated for 20 minutes at 37°C in a thermomixer at 1,100 rpm. After incubation, beads were washed with 1 ml of PNK buffer, then with 1 ml of High-salt buffer, and finally twice with 1 ml of PNK buffer.

### RNA extraction

Beads collected from the previous step were treated with 1 mg/ml Proteinase K (Invitrogen, catalog #AM2544) in 0.25 ml of Proteinase K buffer (100 mM Tris-HCl, pH 7.4; 50 mM NaCl; 10 mM EDTA; 1% SDS) for 20 minutes at 37 °C, shaking at 1,100 rpm. After treatment, 750 μL of Trizol LS (Invitrogen) was added, followed by a total RNA extraction protocol adopted for small RNAs precipiation [2 volumes of Isopropanol instead of 1 with 15 µg of GlycoBlue (Invitrogen, catalog #AM9515); 30 minutes at -80°C; and centrifugation at 18,500 x g for 30-40 minutes at 4°C]. Extracted RNA was stored at -80°C or directly used for RppH treatment.

### RppH treatment

RNA samples were treated with RppH to remove the cap from mRNAs and di- and tri-phosphates from the 5’ end of small RNAs to make them accessible for cloning. RNA extracted from the previous step was treated for one hour at 37°C with 100 μl of RppH reaction mix: 1x NEB Thermopol buffer, RppH (5,000 units/ml) (∼25 units per =< 500 ng of RNA in sample) (NEB, catalog #M0356S). The RppH-treated RNAs were extracted using the Trizol LS (Invitrogen) protocol adopted for small RNA ppt. and stored at -80°C before library preparation.

### RNA libraries and sequencing

Recovered RNAs were used to construct sequencing libraries with the Small RNA-Seq Kit (Lexogen, catalog # 052), which allows simultaneous capture of Argonaute-bound small RNAs and their target RNA fragments (up to ∼200 bp) without separate size-selection steps. Paired-end sequencing (2 × 150 nt) was performed on the Illumina NovaSeq 6000 platform.

### ChIP-seq

Chromatin immunoprecipitation was performed using decoated 4-cell embryos (1.5–3 ml packed embryos per reaction, ∼ 2-4 x 10^7^ embryos). Embryos were resuspended in 20 ml PBS and homogenized with 10 strokes in a metal Dounce homogenizer. Cross-linking was carried out by adding formaldehyde to a final concentration of 2% for 10 minutes at room temperature with gentle rotation, followed by quenching with 0.125 M glycine for 5 minutes on ice. Samples were centrifuged at 780 × g for 5 minutes at 4°C, washed three times with cold PBS, and resuspended in cell lysis buffer (2 mL per 500 μl embryo volume). Chromatin was sheared to a size of 200 – 600 bp using a Bioruptor sonicator (high power, 30 seconds on/off cycles for 15 minutes). Extracts were snap-frozen in liquid nitrogen and stored at -80°C. To verify shearing efficiency, 50 μl of chromatin was reverse cross-linked overnight at 65°C in the presence of 5 M NaCl and Proteinase K, followed by phenol–chloroform extraction and ethanol precipitation, and DNA fragments were analyzed on a 2% agarose gel.

For immunoprecipitation, chromatin extracts were precleared for 2 hours at 4°C with 5 μg of species-matched IgG and 50 μl of 50% Protein A or G agarose slurry (washed three times in TE buffer). Immunoprecipitation was performed overnight at 4°C with 5–20 μg of specific antibody and 50 μl of agarose slurry, followed by sequential 5-minute washes with low-salt, high-salt, LiCl, and TE buffers ^19^. Chromatin was eluted twice with 250 μl elution buffer (1% SDS, 100 mM NaHCO_3_) at room temperature for 15 minutes, and eluates were combined. Cross-links were reversed overnight at 65°C in the presence of 5 M NaCl, EDTA, Tris-HCl (pH 8.0), and Proteinase K (Invitrogen, catalog #AM2544). DNA was purified by phenol-chloroform extraction and ethanol precipitation using glycogen as a carrier and resuspended in 40-80 μl PCR-grade water. The ChIP-seq libraries and sequencing were performed as described ^19^.

### Reads mapping and analysis of DuoRIP-seq

The preprocessing of raw RNA sequencing reads was carried out as previously described ^37^ with modifications (Fig. S10). Briefly, adaptor sequences were trimmed, and length filtering was performed using the FASTX-Toolkit (v0.0.14). The trimmed reads of 18–150 nt were collapsed into non-redundant datasets. To separate siRNA reads from their fragmentated RNA targets, preprocessed 18–150 nt reads were separated into two size fractions: 18–30 nt (small RNAs) and 31–150 nt (long RNA fragments). Then, both fractions were annotated: the 18–30 nt small RNA fraction was annotated by sequential mapping, allowing one mismatch with Bowtie ^65^ to *Ascaris* databases of rRNAs, tRNAs, mitochondrial genome, miRNAs, repetitive elements, and mRNA transcripts, while RNA fragments of 31–150 nt were annotated against the same databases using Bowtie2 ^66^.

Reads were subsequently divided into sense and antisense reads relative to annotated transcripts and intronic or intergenic regions identified. RNA fragments in the 18-30 nt fraction, which lack siRNA signatures, were combined with 31-150 nt, resulting in the 18-150 nt RNA fragments fraction (Fig. S2). For repeat-derived siRNAs, only the 5′ nucleotide information was used due to the lack of strand information for repeats. To avoid ambiguity arising from reads that can be mapped to both repeats and transcripts, sequences were first mapped to repeat elements; the remaining reads were then assigned to mRNA transcripts. Before data normalization, reads corresponding to rRNAs and reads with no genomic match were excluded from the analysis. Expression levels were quantified as counts per million mapped reads (CPM), as this metric directly reflects the relative abundance of Argonaute-bound RNA complexes, where each siRNA read typically corresponds to a single target RNA fragment. The Bowtie2 mapping result was used to determine the expression levels of corresponding targets.

Only genomic regions with ≥6× CPM read coverage were considered for downstream analysis. The average of 2-3 biological replicates from each sample was used, and a two-fold expression difference was applied as a threshold for differential expression. Comparative analyses were conducted using a custom pipeline that uses BEDTools ^67^ and SAMTools ^68^.

### Repetitive element definition and assessment of multimapping behavior

Repetitive elements (hereafter referred to as “repeats”) were defined based on our previous study ^37^. Briefly, genomic regions exhibiting ≥10-fold enrichment of WAGO-associated 22G-RNAs (AsWAGO-1, AsWAGO-2, and AsNRDE-3) in at least one dataset, after excluding rRNA, tRNA, miRNA, and mitochondrial sequences, were classified as WAGO-repeats. These regions represent loci enriched for endogenous siRNAs and were used as the reference WAGO-siRNA-targeted repeats (WAGO-repeats) in this study. Thus, WAGO-repeats represent a functional subset of genomic regions enriched for WAGO associated endogenous siRNA rather than all repetitive DNA.

Because repeat-derived siRNAs can map to multiple genomic loci, we evaluated whether multimapping could influence the observed repeat-associated enrichment patterns. Primary siRNA analysis used the standard Bowtie2 alignment strategy described above, in which a single best-scoring alignment was reported for each read. An independent ambiguity-space analysis was performed to assess the potential impact of this assignment strategy. For this analysis Bowtie2 alignment were generated while retaining all equivalent best-scoring genomic alignments. Reads with multiple equal-best alignments were grouped into ambiguity sets defined by their collection of equivalent genomic mapping locations. Rarefaction and site-occupancy analyses were used to assess the reuse and sampling saturation of these ambiguity sets and their corresponding genomic sites with increasing sequencing depth (Fig. S9).

This analysis was performed to evaluate the robustness of repeat-associated enrichment to multimapping. Because equivalent alignments cannot establish the specific repeat copy from which an individual siRNA originated, the analysis does not resolve copy-level attribution. Rather, it tests whether the observed repeat-associated signal is supported by recurrently sampled ambiguity structures or could be explained predominantly by sparse, stochastic multimapping. This analysis showed that the observed recurrence and saturation of ambiguity structures with increasing sequencing depth support the robustness of the repeat-associated enrichment patterns (Fig. S9).

### ChIP-seq analysis

Sequencing reads from ChIP-seq experiments were aligned to the reference genome using Bowtie2 ^66^, and duplicate reads were removed before peak detection. Enrichment peaks were identified using MACS3 ^69^ (*v*3.0.1; Model-based Analysis of ChIP-Seq) with default settings. Broad peaks were used to define significant binding regions and their corresponding summit positions for downstream analyses. Only peaks passing the experiment-specific significance threshold (q < 0.05) were retained for further evaluation.

To improve reproducibility across biological replicates, we adopted a strategy that combines replicate information based on peak significance (p-values) rather than requiring peak presence in all replicates. Peaks were first called independently for each replicate using MACS3, and the resulting peak sets were subsequently processed with MSPC (Multiple Sample Peak Calling). MSPC combines p-values across replicates, applies a right-tail probability threshold to retain only statistically reproducible peaks, and generates a consensus set of high-confidence binding regions for downstream analyses.

### Data visualization and statistical analysis

All plots were generated using custom scripts in R (*ggplot2*) and Python (*matplotlib*). Circular histogram plots were produced using Circos (v0.69-9) ^70^. Genome browser tracks were generated in BigWig format for visualization in the UCSC Genome Browser ^71^.

## Data and resource availability

The sequencing data are deposited in the NCBI GEO database with accession number GSE316157. The data are also available in UCSC Genome Browser track data hubs that can be accessed with this link: https://genome.ucsc.edu/s/MVZ/small_RNAs.

## Author contributions

R.E.D. and J.W. conceived the study; M.V.Z. performed the WAGO-IP, cellular fractionation, and DuoRIP-seq; M.V.Z. carried out bioinformatics analysis; Y.K. did ChIP-seq; J.W. performed immunohistochemistry; M.V.Z. wrote the initial manuscript draft; M.V.Z., R.E.D., and J.W. edited the manuscript; and R.E.D. and J.W. provided supervision, project administration, and funding.

## Acknowledgements

We thank Bruce Bamber, Jeff Myers, and Routh Packing Co. for their support and hospitality in collecting *Ascaris* material. We thank Adam Wallace and Stella Kratzer for initial work on the purification, evaluation, and Western blot analysis of the *Ascaris* Argonaute antibodies. We thank Ashley Neff for the preliminary IHC staining and exploratory efforts examining small RNAs associated with Argonaute antibodies.

## Supplementary Figure Legends

**Figure S1. Additional staining of WAGOs in *Ascaris* embryos.** Immunohistochemistry staining of early *Ascaris* embryos using antibodies against *Ascaris* Argonaute proteins AsWAGO-1, AsWAGO-2, AsWAGO-3, AsCSR-1, and AsNRDE-3. Elimination and non-elimination anaphase mitoses are shown, highlighting the localization of AsWAGO-2 and AsWAGO-3 to chromatin undergoing programmed DNA elimination.

**Figure S2. Bioinformatic separation of antisense siRNAs (18-30 nt) and target RNAs (18-150 nt) after RNase treatment.** Annotation categories: exons, introns, WAGO-repeats, and intergenic. Colored bars indicate the criteria used for read separation, classifying reads to either siRNAs, mRNAs, or indicating the presence of dsRNA where the strand information can be identified.

**Figure S3. RNA expression profiles of *Ascaris* Argonautes WAGO-2, WAGO-3, CSR-1, and WAGO-1 throughout the early development.** The Argonautes are plotted in two groups based on their expression level. Embryogenesis stages E1-E10: zygote maturation stages prior to pronuclear fusion isolated from the uterus (E1–4; E4 is the stage passed from Ascaris and the host to the environment); embryo development (at 30 °C) stages, with E5=24 h (1-cell), E6=46 h (2-cell), E7=64 h (4-cell), E8 =96 h (16-cell), E9 = 116 h (32–64-cell), E10 = 7day (256-cell), larvae L1 (10-day) and L2 (21-day). The RNA-seq expression data are provided as a Source Data file.

**Figure S4. *Ascaris* Argonautes are associated with specific small RNAs and their targets.** Additional size distribution plots and targets for all WAGO IPs for biological replicates.

**Figure S5. AsWAGO-2 and AsWAGO-3 IP in subcellular fractionation captures siRNA and target RNAs.** Unique and overlapping AsWAGO-2 and AsWAGO-3 genes and repeats associated RNA targets and siRNA among the three subcellular fractions (Left). Genome-wide comparison of the nuclear fractions (chromatin and nucleoplasm, x-axis) with cytoplasm (y-axis) for the repeats and genes, siRNA, and target RNAs on the dense scatter plots.

**Figure S6. AsWAGO-3 associated small RNAs predominantly target DNA repeats in the middle of the chromosomes that are eliminated by PDE.** “Right” eliminated regions. AsWAGO-3 associated small RNAs and target RNAs from the chromatin fraction mapped to the right (R) eliminated regions (a similar pattern was observed for the left elimination region) (see Figure 5 for the data and the legend).

**Figure S7. AsWAGO-3-associated small RNAs predominantly target eliminated DNA repeats in the middle of the chromosomes (nucleoplasm and cytoplasm fractions).** Levels of AsWAGO-3-associated siRNAs and target RNAs (CPM, square root scale) in eliminated regions are shown in the raincloud plots (legend as Figure 5C). siRNAs targeting repeats are significantly higher in the middle (M, black *) than at the ends (L/R) of the chromosomes. In contrast to the chromatin, the regions eliminated in the middle of chromosomes do not show a higher level of target RNA transcription (L/R is significantly higher, black *) in nucleoplasm and cytoplasm fractions. Similar to the chromatin pattern, siRNAs targeting genes are more enriched at the ends (L/R, red *) than in the middle (M) of the chromosomes. Regions eliminated at the ends of chromosomes show much higher levels of target RNAs (L/R, green *). The data are shown in a square root scale of CPM. Wilcoxon test (two-tailed) was used for significance; effect sizes are given as Cliff’s delta.

**Figure S8. AsWAGO-2 and AsWAGO-3-associated siRNAs and target RNAs across cellular fractions**. Enrichment of siRNAs and target RNAs in the nucleus for AsWAGO-2 **(A)** and AsWAGO-3 **(B)**. The abundance of AsWAGO-associated siRNAs (top) and RNAs (bottom) mapped to eliminated (left) and retained (right) genomic regions (in CPM, log₁₀ scale). Red and blue points represent eliminated and retained loci, respectively; green marks siRNAs or RNAs associated with enriched fraction AsWAGO siRNAs targets that have >2-fold enrichment in nuclear fraction (X-axis).

**Figure S9. Evaluation of multimapping of repeat-associated siRNAs using rarefaction and saturation analyses.** Rarefaction analyses were performed using Bowtie2 alignments retaining all equivalent best-scoring genomic alignments to characterize sampling of the multimapping ambiguity space. Reads were randomly subsampled at increasing fractions of the dataset (1%, 2%, 5%, 10%, 20%, 40%, 60%, 80%, and 100%), with five independent random subsamples performed at each depth. **A. Ambiguity-set rarefaction** shows the number of unique ambiguity sets identified as a function of the number of sampled reads. **B. Good’s coverage** estimates the proportion of the observed ambiguity-set space represented by non-singleton observations; increasing values and a plateau indicate diminishing discovery of previously unobserved ambiguity sets with additional sequencing. **C. Discovery decay** shows the number of newly identified ambiguity sets per additional sampled read, providing a direct measure of diminishing discovery with increasing sequencing depth. **D. Singleton decay** shows the fraction of ambiguity sets represented by a single read at each sampling depth. **E. Ambiguity-set reuse saturation** shows the mean number of reads assigned per observed ambiguity set, reflecting increasing reuse of recurrent multimapping structures with sequencing depth. Lines represent the mean of five independent subsamples.

**Figure S10. Bioinformatic workflow for processing and analysis of small RNA and target RNA sequencing data.** The flowchart summarizes the sequential processing, annotation, classification, genomic remapping, filtering, and quantification of sequencing reads. Standard flowchart symbols indicate different types of operations: **rounded rectangles** represent processes or computational steps, **diamonds** represent decision points or classification criteria, **parallelograms** represent input/output data, and **dashed boxes** indicate groups of related classifications or outputs. Arrows indicate the direction of data processing. Read classification incorporates read length, strand orientation, 5′ nucleotide identity, and genomic annotation to distinguish siRNAs, target RNA fragments, and dsRNA-associated signals. Genome-mapped reads are subsequently filtered, normalized, quantified, and combined for downstream analyses.

