## Supplemental Figures for "Argonautes and small RNAs associated with nematode programmed DNA elimination"

Fig. S1

### AsWAGO-3

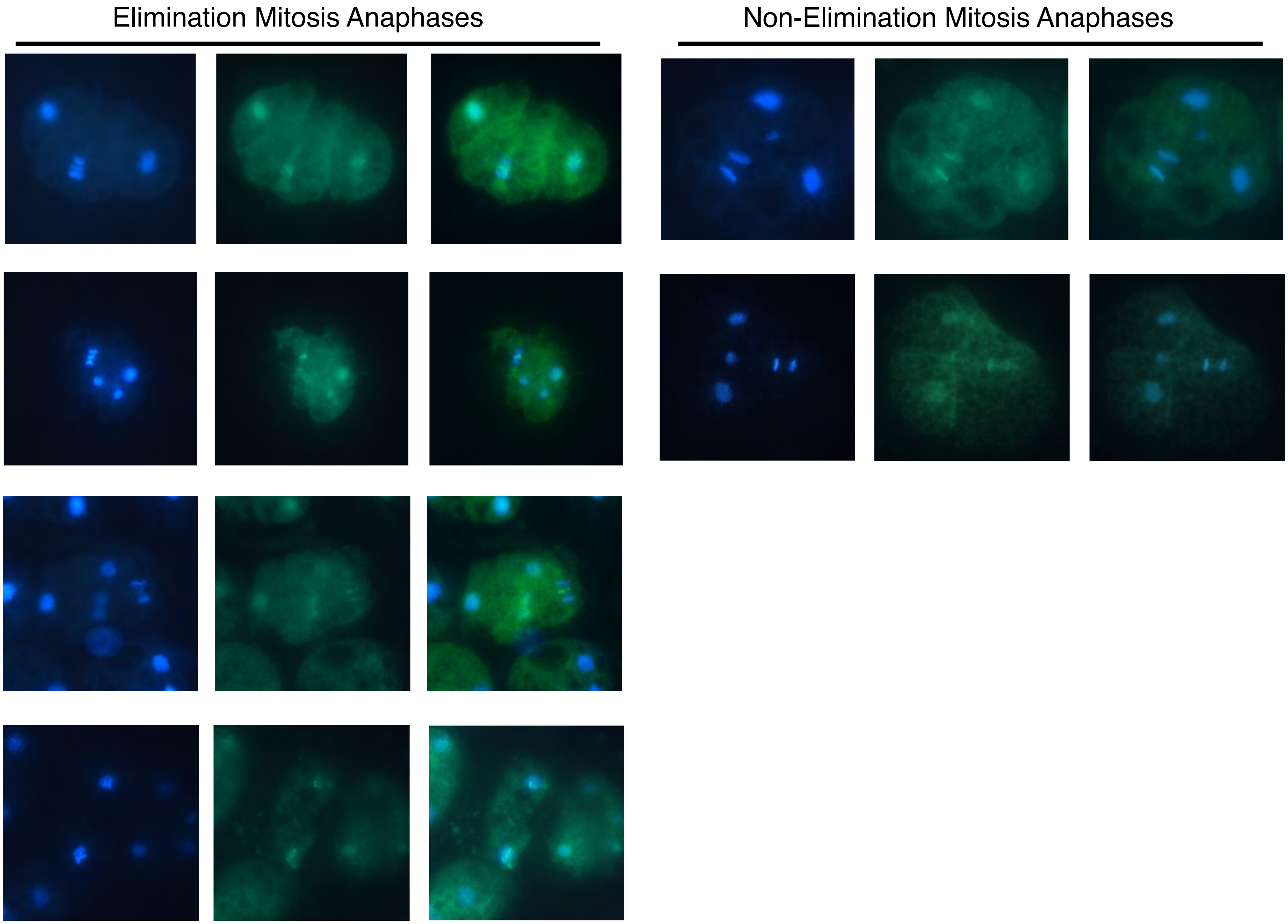

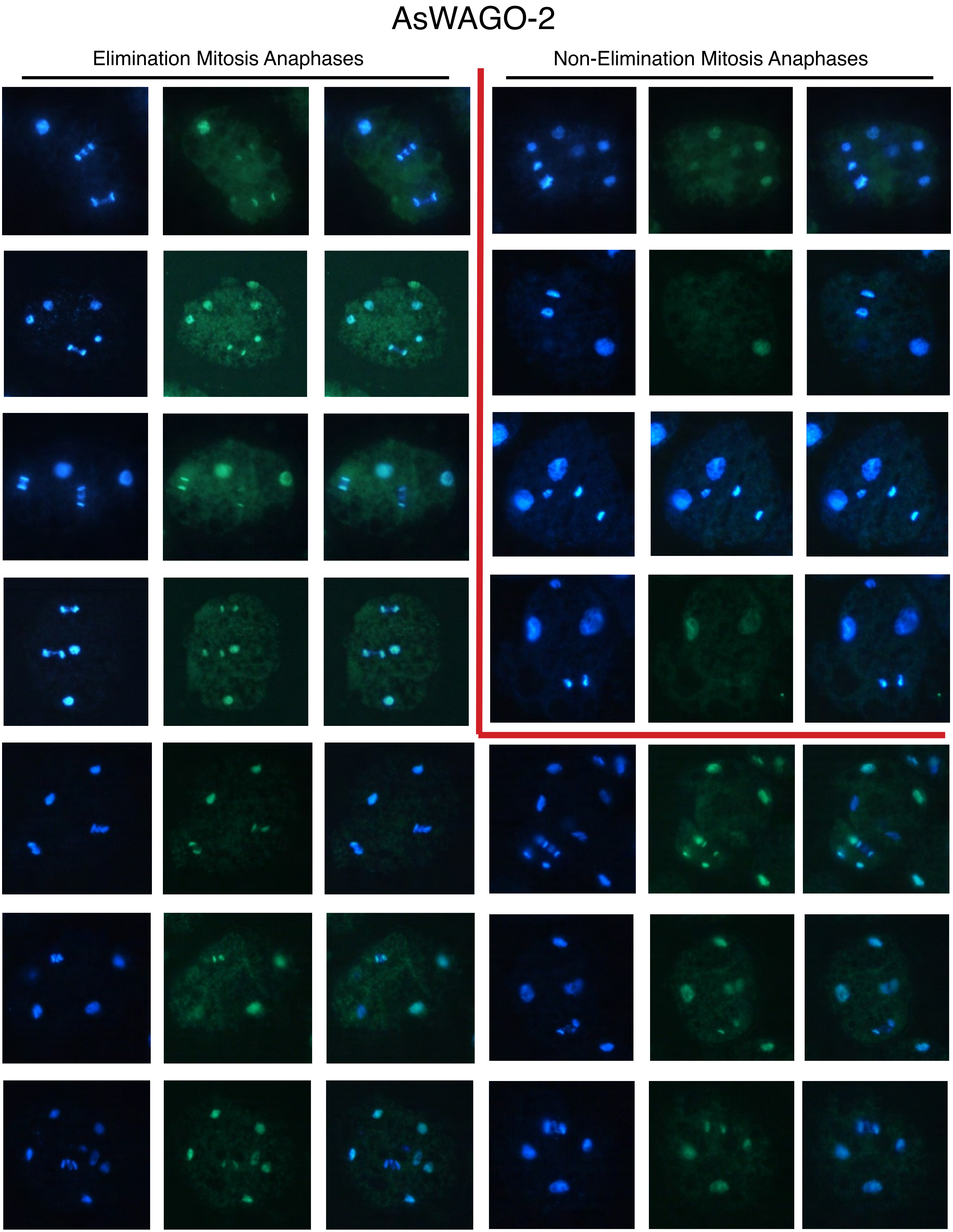
AsWAGO-2
Elimination Mitosis Anaphases
Non-Elimination Mitosis Anaphases

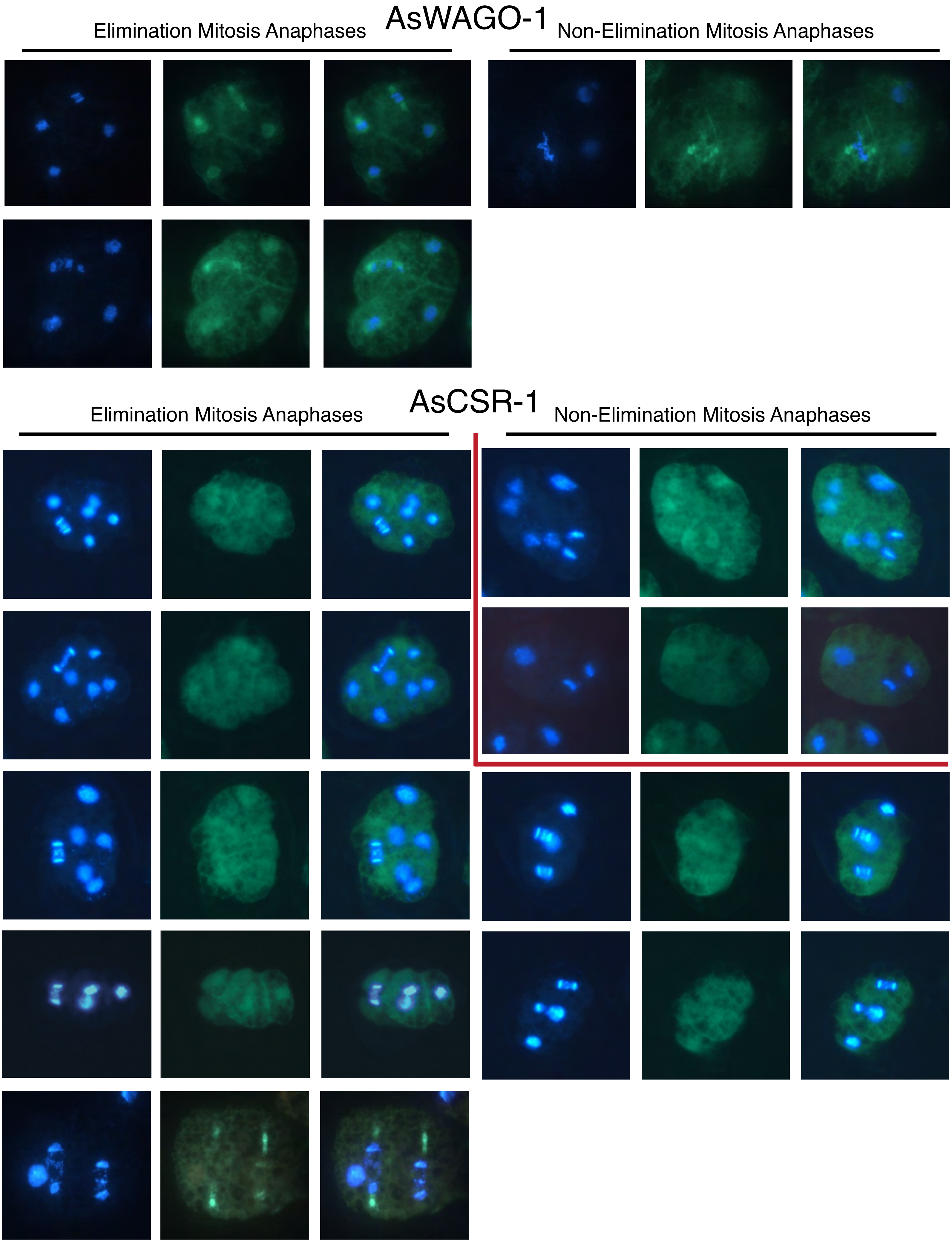
AsWAGO-1
Elimination Mitosis Anaphases
Non-Elimination Mitosis Anaphases
AsCSR-1
Elimination Mitosis Anaphases
Non-Elimination Mitosis Anaphases

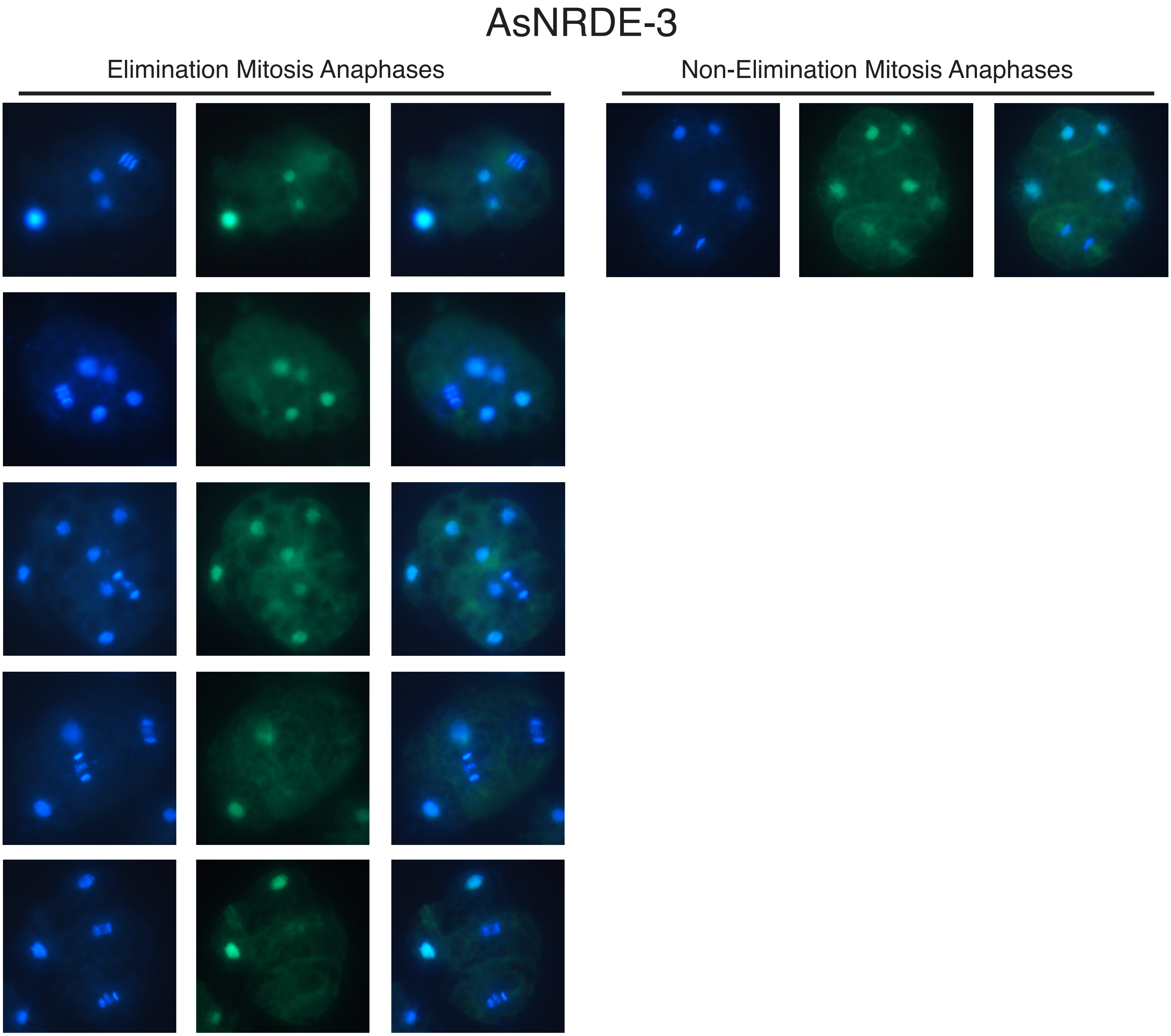
AsNRDE-3
Elimination Mitosis Anaphases
Non-Elimination Mitosis Anaphases

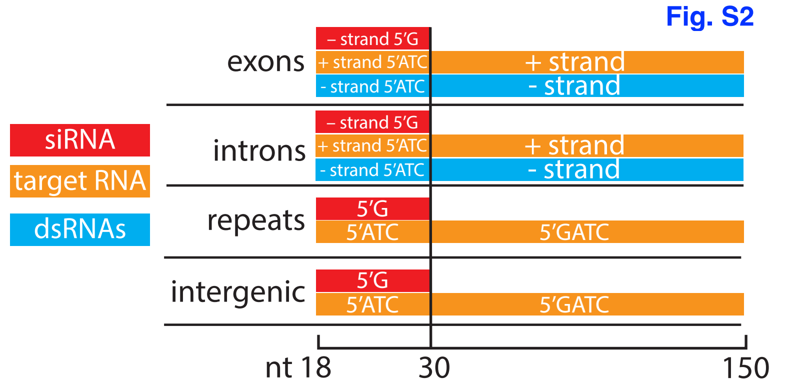
Fig. S2
siRNA
target RNA
dsRNAs
exons
– strand 5′G
+ strand 5′ATC
- strand 5′ATC
+ strand
- strand
introns
– strand 5′G
+ strand 5′ATC
- strand 5′ATC
+ strand
- strand
repeats
5′G
5′ATC
5′GATC
intergenic
5′G
5′ATC
5′GATC
nt 18
30
150

Fig. S3

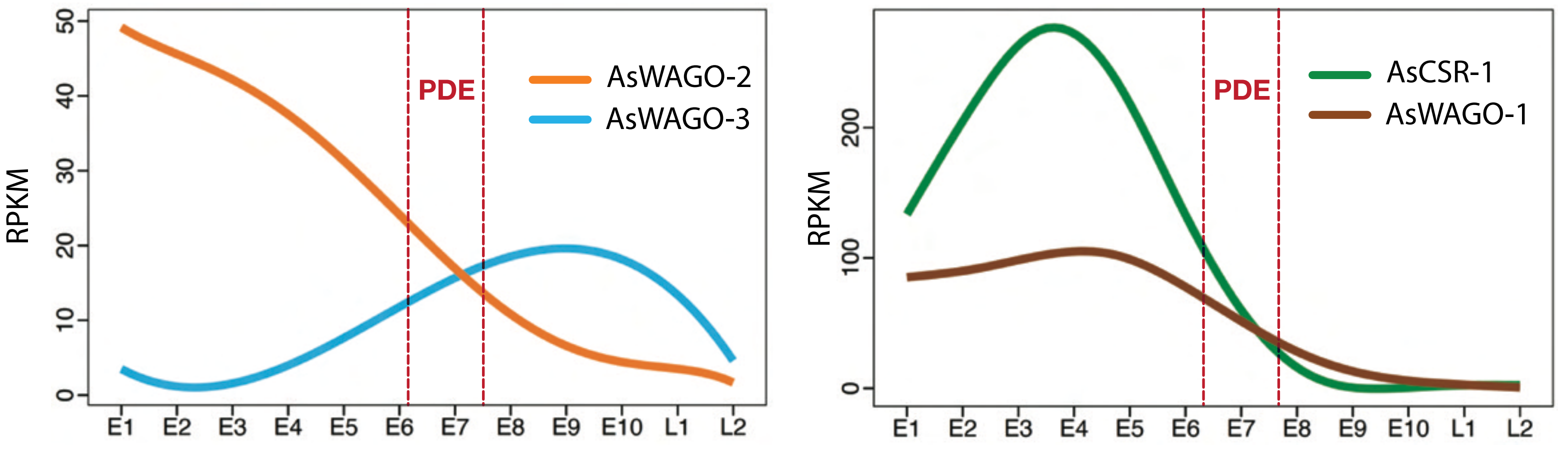

RPKM
50
40
30
20
10
0
PDE
AsWAGO-2
AsWAGO-3
E1
E2
E3
E4
E5
E6
E7
E8
E9
E10
L1
L2
RPKM
200
100
0
PDE
AsCSR-1
AsWAGO-1
E1
E2
E3
E4
E5
E6
E7
E8
E9
E10
L1
L2

Fig. S4

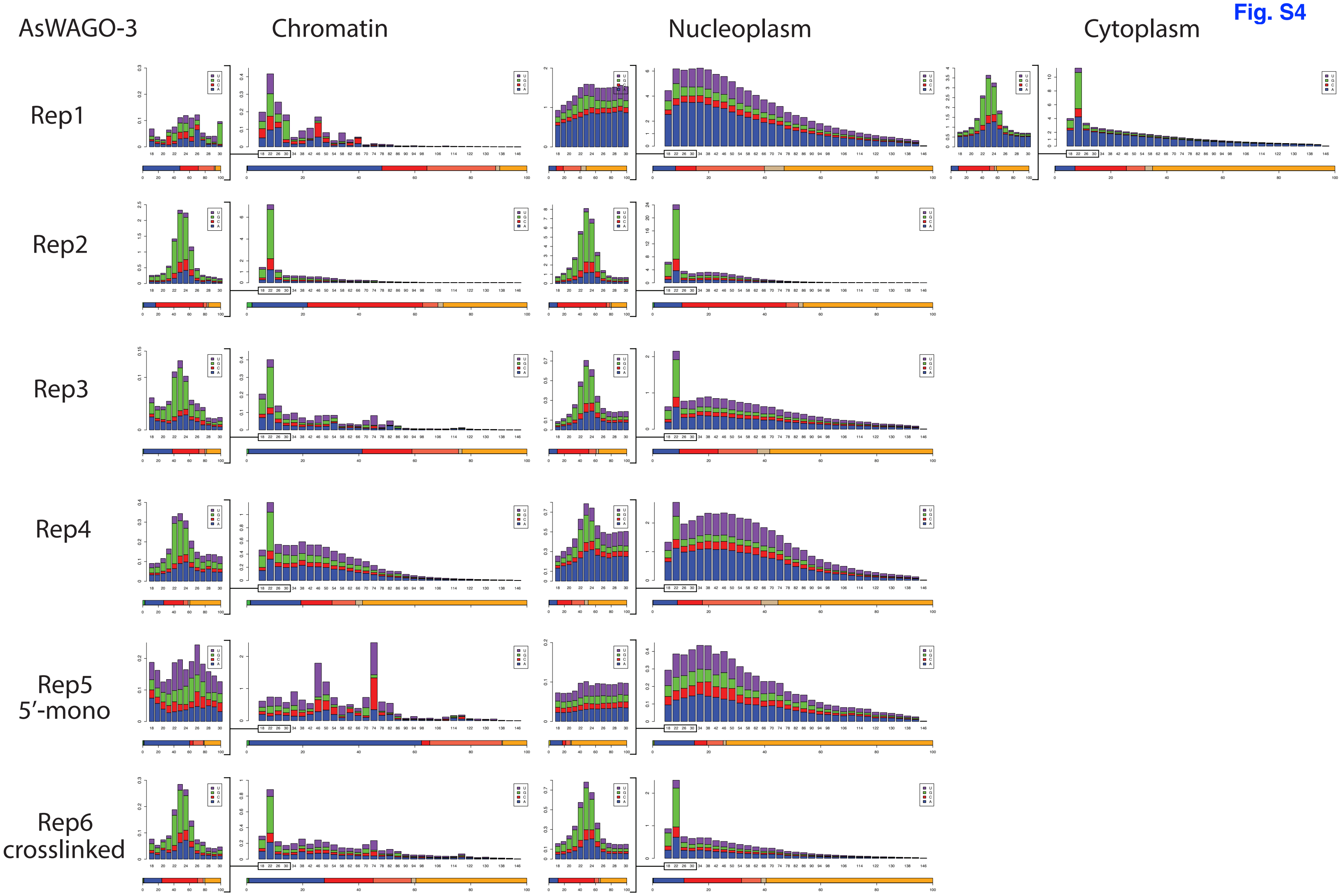

AsWAGO-3
Chromatin
Nucleoplasm
Cytoplasm
Rep1
Rep2
Rep3
Rep4
Rep5
5′-mono
Rep6
crosslinked

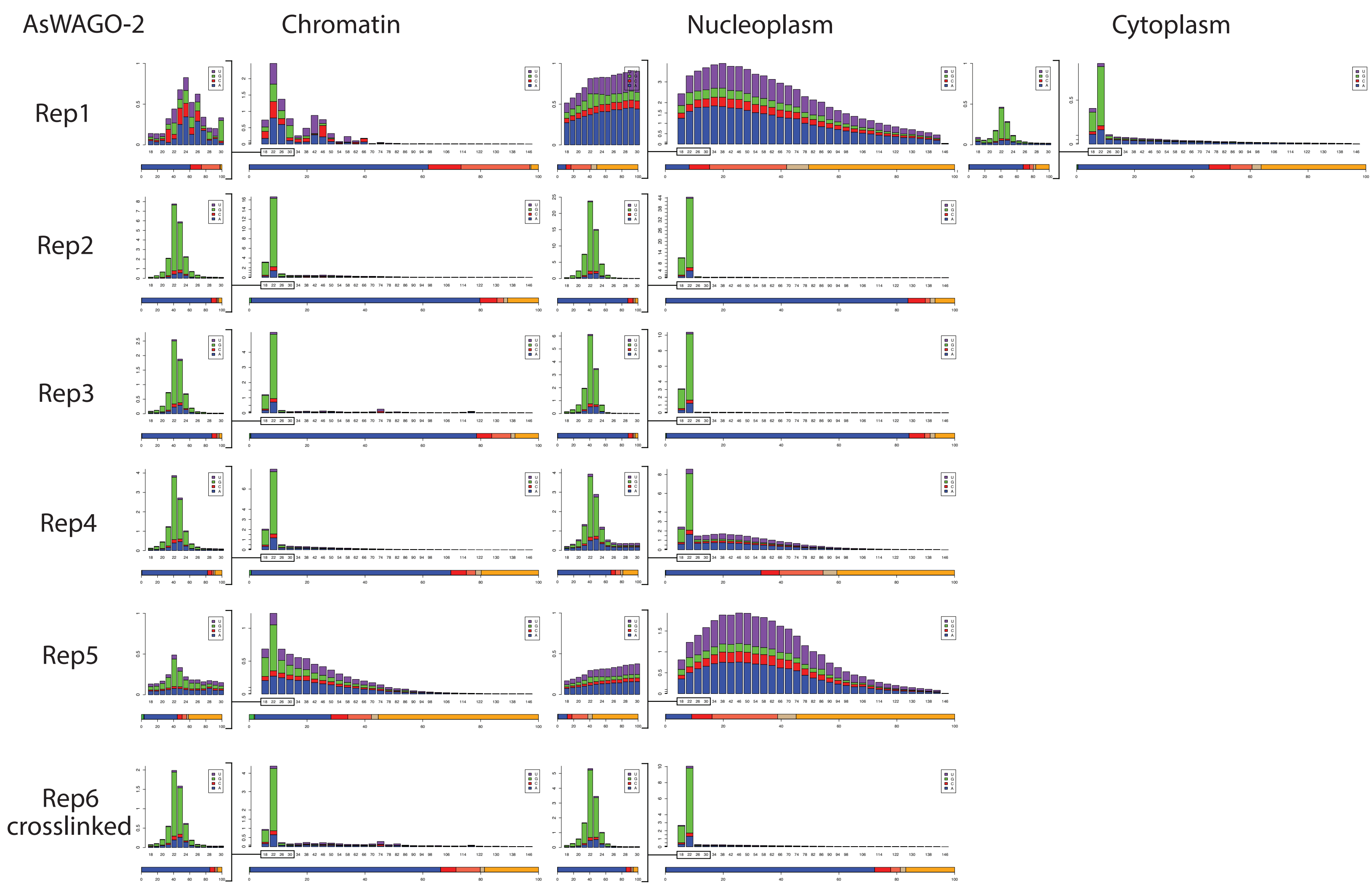

AsWAGO-2
Chromatin
Nucleoplasm
Cytoplasm
Rep1
Rep2
Rep3
Rep4
Rep5
Rep6
crosslinked

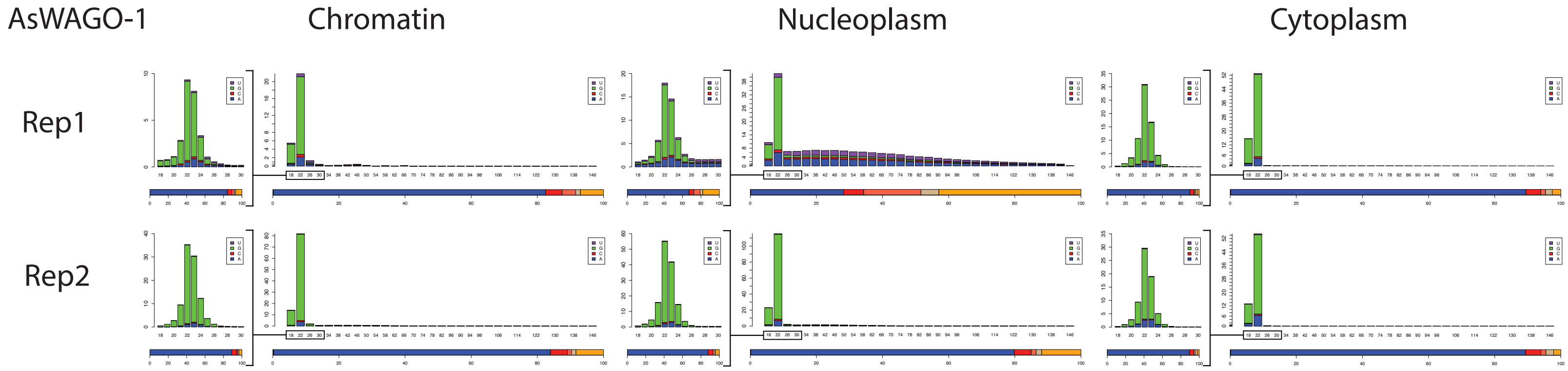
AsWAGO-1
Chromatin
Nucleoplasm
Cytoplasm
Rep1
Rep2

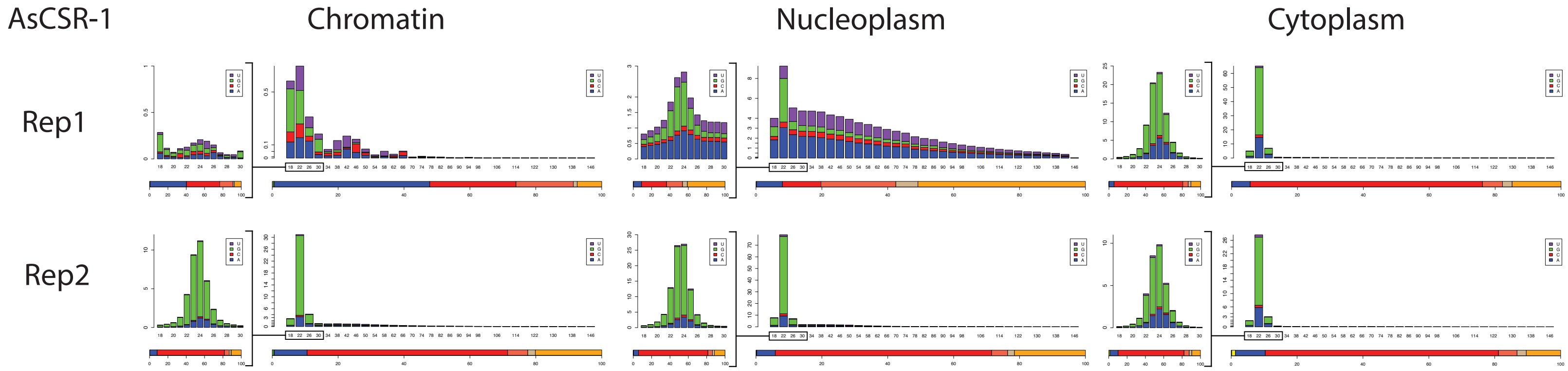

AsCSR-1
Chromatin
Nucleoplasm
Cytoplasm
Rep1
Rep2

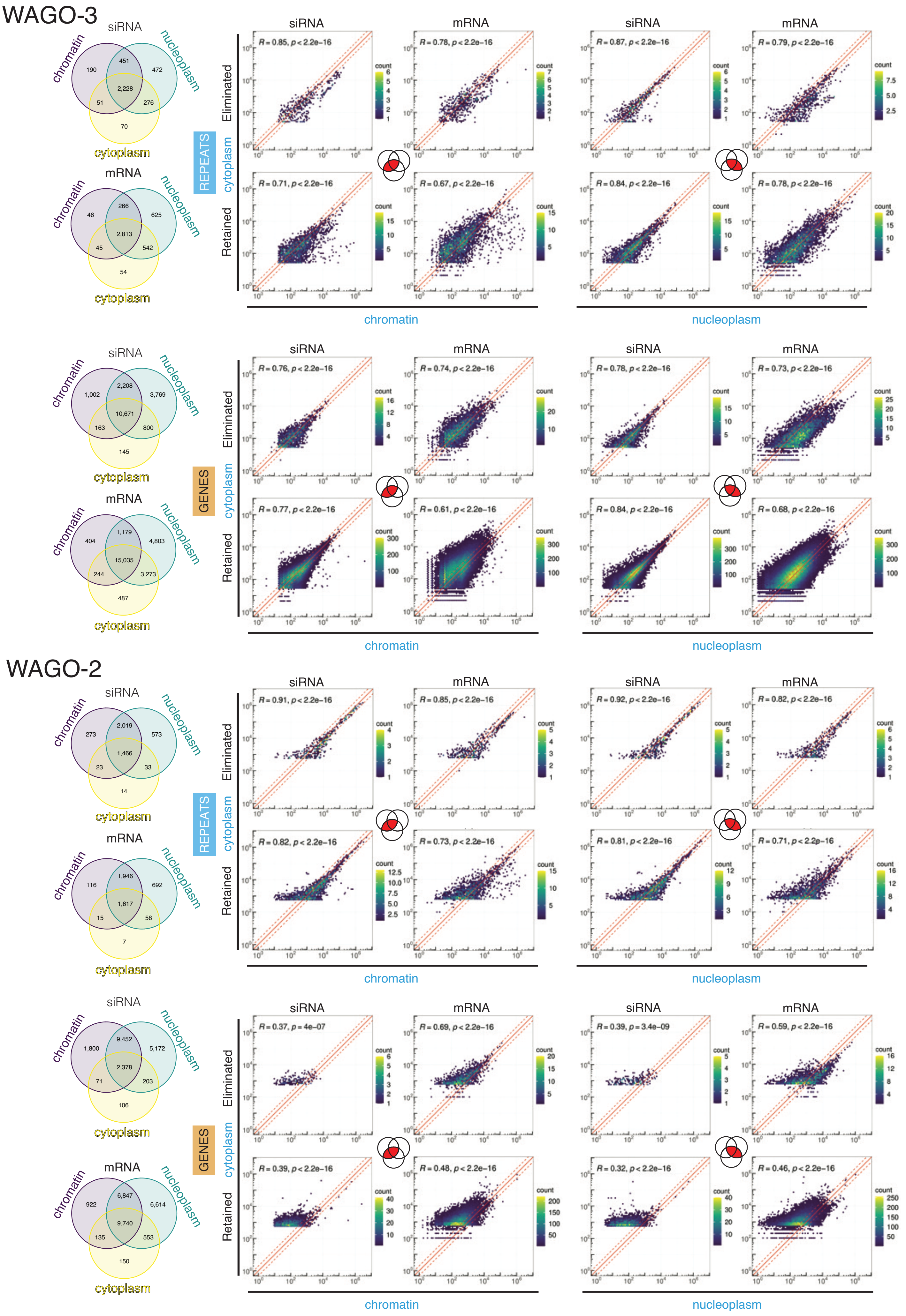

WAGO-3
REPEATS
siRNA
chromatin 190 451 472 nucleoplasm
2,228
51 276
70
cytoplasm
mRNA
chromatin 46 266 625 nucleoplasm
2,813
45 542
54
cytoplasm
Eliminated
Retained
cytoplasm
siRNA mRNA siRNA mRNA
R = 0.85, p < 2.2e−16
R = 0.78, p < 2.2e−16
R = 0.87, p < 2.2e−16
R = 0.79, p < 2.2e−16
R = 0.71, p < 2.2e−16
R = 0.67, p < 2.2e−16
R = 0.84, p < 2.2e−16
R = 0.78, p < 2.2e−16
count
chromatin
nucleoplasm
GENES
siRNA
chromatin 1,002 2,208 3,769 nucleoplasm
10,671
163 800
145
cytoplasm
mRNA
chromatin 404 1,179 4,803 nucleoplasm
15,035
244 3,273
487
cytoplasm
R = 0.76, p < 2.2e−16
R = 0.74, p < 2.2e−16
R = 0.78, p < 2.2e−16
R = 0.73, p < 2.2e−16
R = 0.77, p < 2.2e−16
R = 0.61, p < 2.2e−16
R = 0.84, p < 2.2e−16
R = 0.68, p < 2.2e−16
WAGO-2
REPEATS
siRNA
chromatin 273 2,019 573 nucleoplasm
1,466
23 33
14
cytoplasm
mRNA
chromatin 116 1,946 692 nucleoplasm
1,617
15 58
7
cytoplasm
R = 0.91, p < 2.2e−16
R = 0.85, p < 2.2e−16
R = 0.92, p < 2.2e−16
R = 0.82, p < 2.2e−16
R = 0.82, p < 2.2e−16
R = 0.73, p < 2.2e−16
R = 0.81, p < 2.2e−16
R = 0.71, p < 2.2e−16
GENES
siRNA
chromatin 1,800 9,452 5,172 nucleoplasm
2,378
71 203
106
cytoplasm
mRNA
chromatin 922 6,847 6,614 nucleoplasm
9,740
135 553
150
cytoplasm
R = 0.37, p = 4e−07
R = 0.69, p < 2.2e−16
R = 0.39, p = 3.4e−09
R = 0.59, p < 2.2e−16
R = 0.39, p < 2.2e−16
R = 0.48, p < 2.2e−16
R = 0.32, p < 2.2e−16
R = 0.46, p < 2.2e−16


Fig. S5

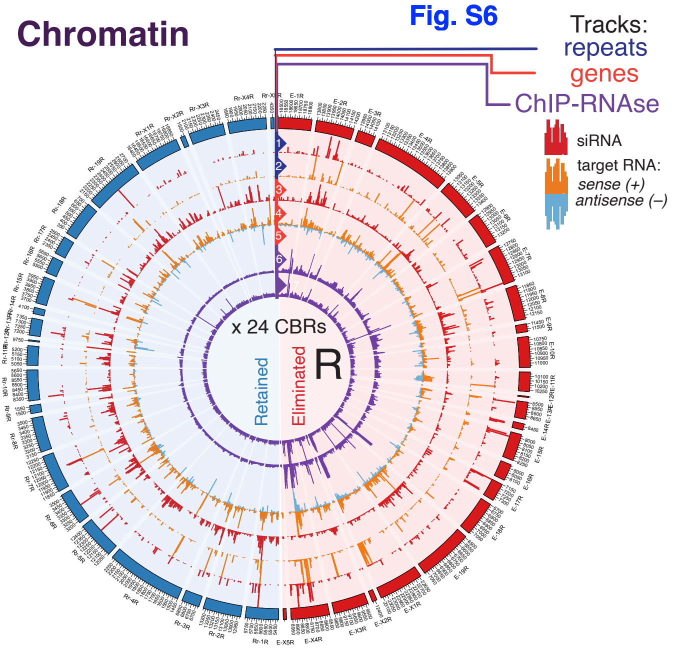
Chromatin
Fig. S6
Tracks:
repeats
genes
ChIP-RNAse
siRNA
target RNA:
sense (+)
antisense (–)
x 24 CBRs
Retained
Eliminated
R

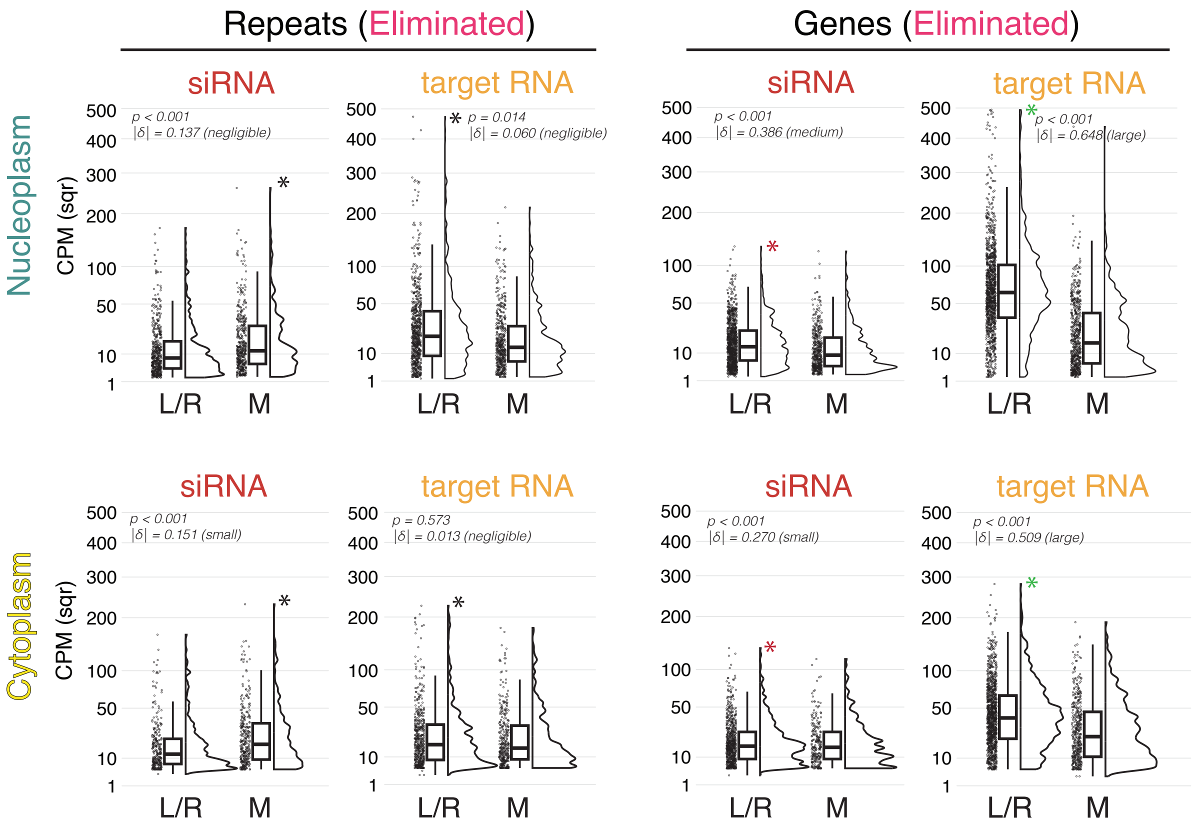
Repeats (Eliminated)
Genes (Eliminated)
Nucleoplasm
Cytoplasm
siRNA
target RNA
CPM (sqr)
p < 0.001
|δ| = 0.137 (negligible)
p = 0.014
|δ| = 0.060 (negligible)
p < 0.001
|δ| = 0.386 (medium)
p < 0.001
|δ| = 0.648 (large)
p < 0.001
|δ| = 0.151 (small)
p = 0.573
|δ| = 0.013 (negligible)
p < 0.001
|δ| = 0.270 (small)
p < 0.001
|δ| = 0.509 (large)
L/R
M

Fig. S8 A

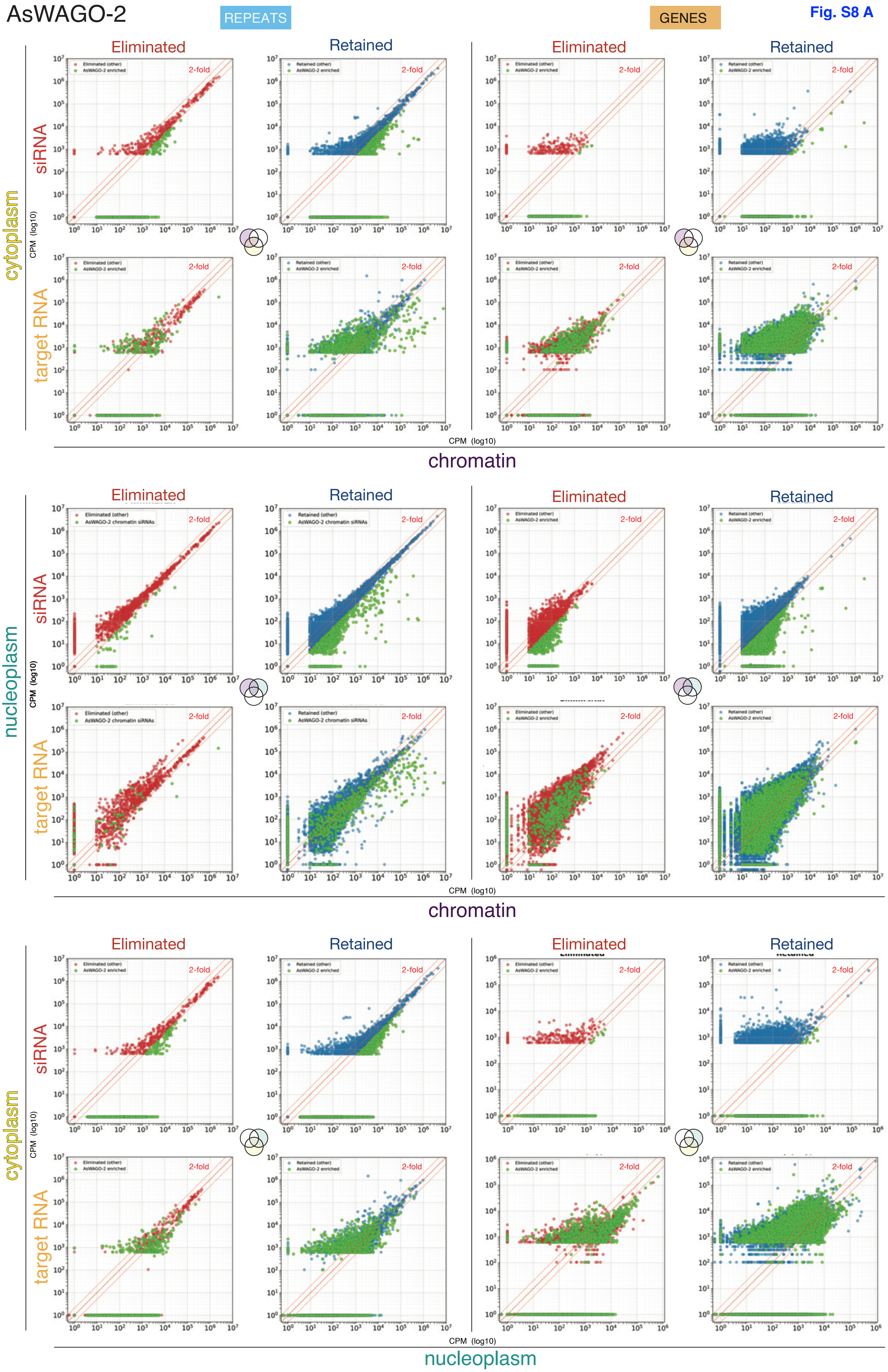
AsWAGO-2
REPEATS
GENES
Eliminated
Retained
siRNA
target RNA
cytoplasm
nucleoplasm
CPM (log10)
chromatin
2-fold
Eliminated (other)
Retained (other)
AsWAGO-2 enriched
AsWAGO-2 chromatin siRNAs

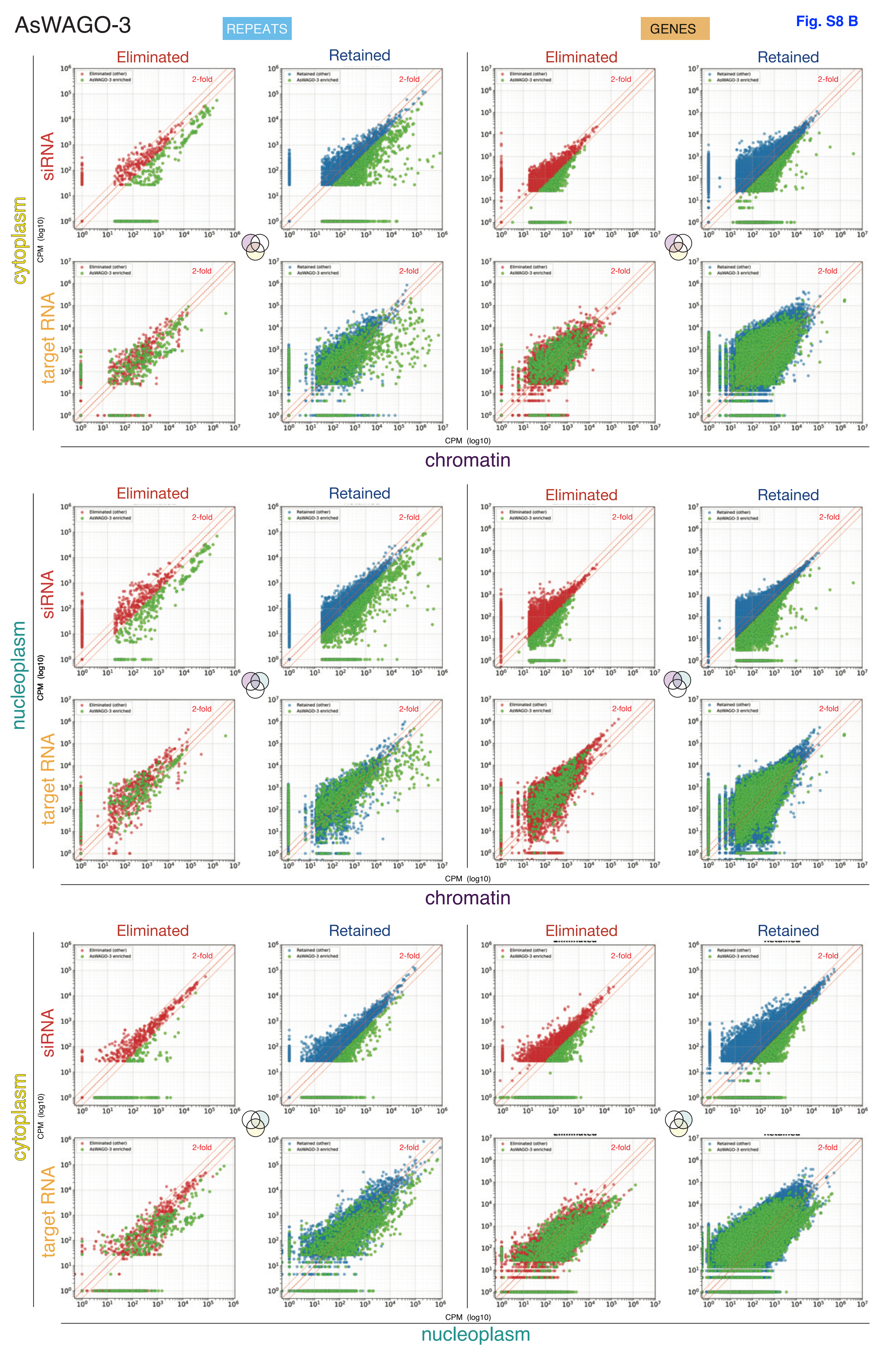

AsWAGO-3
REPEATS
GENES
Fig. S8 B
Eliminated
Retained
siRNA
target RNA
cytoplasm
nucleoplasm
chromatin
CPM (log10)
2-fold
Eliminated (other)
Retained (other)
AsWAGO-3 enriched

Fig. S9

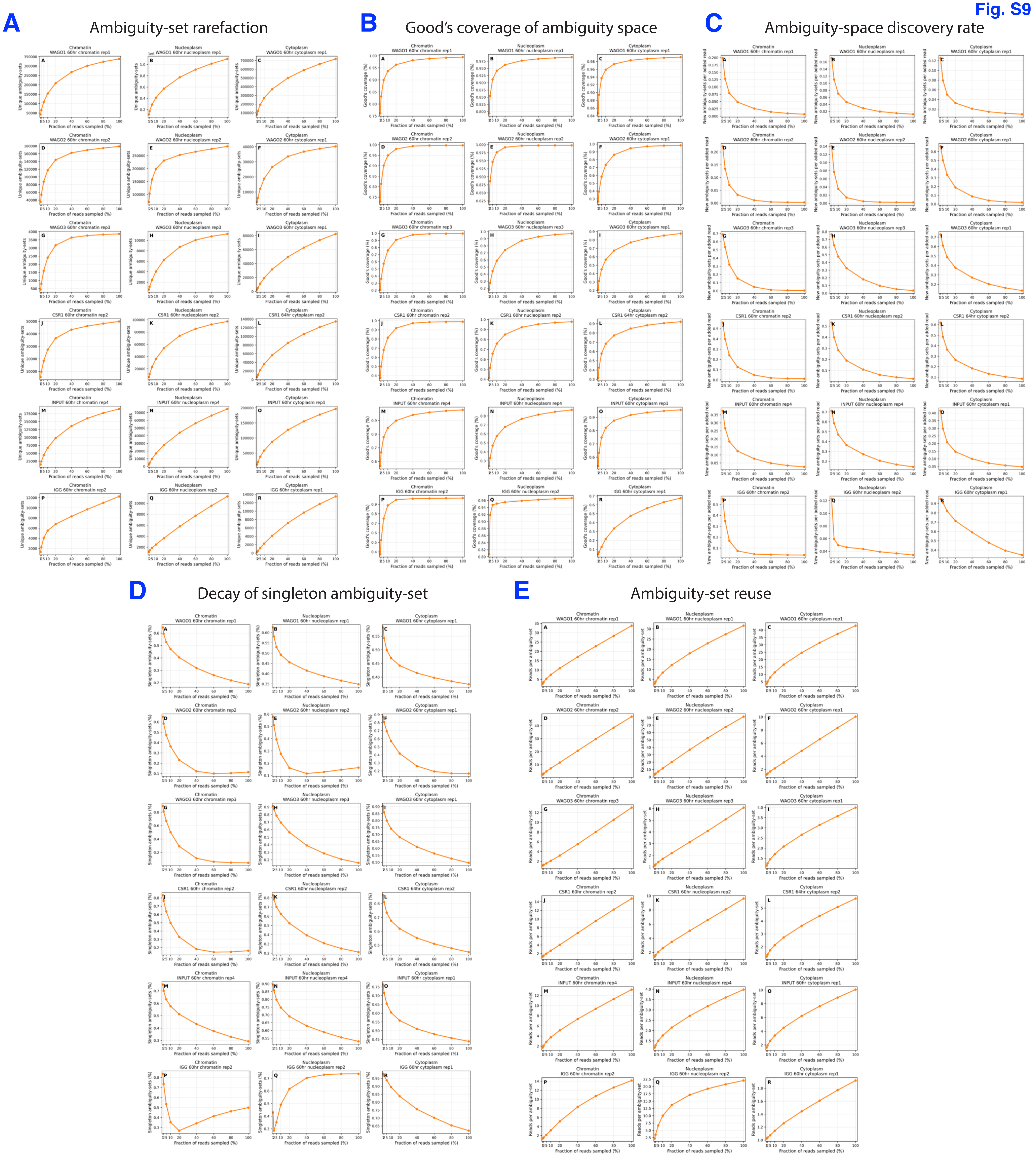
A
Ambiguity-set rarefaction
B
Good's coverage of ambiguity space
C
Ambiguity-space discovery rate
D
Decay of singleton ambiguity-set
E
Ambiguity-set reuse

Fig. S10

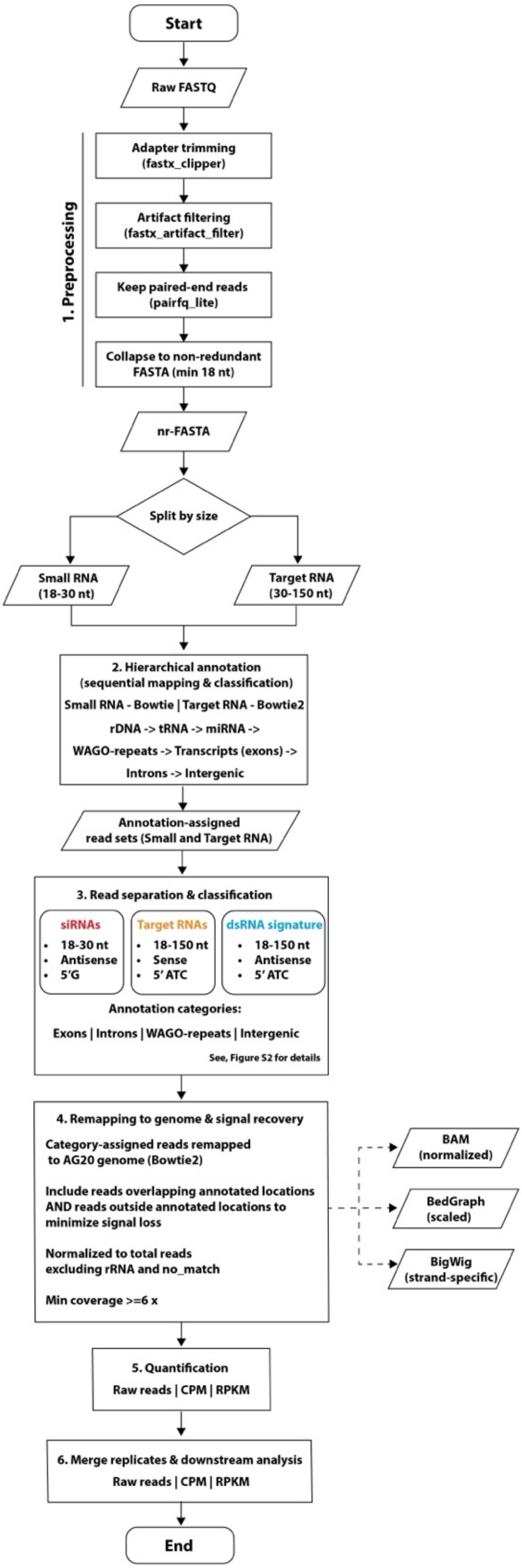
Start
Raw FASTQ
1. Preprocessing
Adapter trimming (fastx_clipper)
Artifact filtering (fastx_artifact_filter)
Keep paired-end reads (pairfq_lite)
Collapse to non-redundant FASTA (min 18 nt)
nr-FASTA
Split by size
Small RNA (18-30 nt)
Target RNA (30-150 nt)
2. Hierarchical annotation (sequential mapping & classification)
Small RNA - Bowtie | Target RNA - Bowtie2
rDNA -> tRNA -> miRNA ->
WAGO-repeats -> Transcripts (exons) ->
Introns -> Intergenic
Annotation-assigned read sets (Small and Target RNA)
3. Read separation & classification
siRNAs
18-30 nt
Antisense
5'G
Target RNAs
18-150 nt
Sense
5' ATC
dsRNA signature
18-150 nt
Antisense
5' ATC
Annotation categories:
Exons | Introns | WAGO-repeats | Intergenic
See, Figure S2 for details
4. Remapping to genome & signal recovery
Category-assigned reads remapped to AG20 genome (Bowtie2)
Include reads overlapping annotated locations AND reads outside annotated locations to minimize signal loss
Normalized to total reads excluding rRNA and no_match
Min coverage >=6 x
BAM (normalized)
BedGraph (scaled)
BigWig (strand-specific)
5. Quantification
Raw reads | CPM | RPKM
6. Merge replicates & downstream analysis
Raw reads | CPM | RPKM
End
